# Hierarchical Interleaved Bloom Filter: Enabling ultrafast, approximate sequence queries

**DOI:** 10.1101/2022.08.01.502266

**Authors:** Svenja Mehringer, Enrico Seiler, Felix Droop, Mitra Darvish, René Rahn, Martin Vingron, Knut Reinert

**Affiliations:** Department of Mathematics and Computer Science, Freie Universität Berlin, Takustr. 9, 14195 Berlin, Germany; MPI for Molecular Genetics, Ihnestr. 63, 14195 Berlin, Germany

## Abstract

Searching sequences in large, distributed databases is the most widely used bioinformatics analysis done. This basic task is in dire need for solutions that deal with the exponential growth of sequence repositories and perform approximate queries very fast.

In this paper, we present a novel data structure: the Hierarchical Interleaved Bloom Filter (HIBF). It is extremely fast and space efficient, yet so general that it has the potential to serve as the underlying engine for many applications.

We show that the HIBF is superior in build time, index size and search time while achieving a comparable or better accuracy compared to other state-of-the art tools (Mantis and Bifrost). The HIBF builds an index up to 211 times faster, using up to 14 times less space and can answer approximate membership queries faster by a factor of up to 129. This can be considered a quantum leap that opens the door to indexing complete sequence archives like the European Nucleotide Archive or even larger metagenomics data sets.

## 1 Introduction

Following the sequencing of the human genome [VR^+^01, Int01], genomic analysis has come a long way. The recent improvements of sequencing technologies, commonly subsumed under the term NGS (Next-Generation Sequencing) or 3rd (and 4th) generation sequencing, have triggered incredible innovative diagnosis and treatments in biomedicine, but also tremendously increased the sequencing throughput. Within 10 years, the current throughput of standard Illumina machines rose from 21 billion base pairs [VR^+^01, Int01] collected over months to about 3000 billion base pairs per day.

As a result of this development, the number of new data submissions, generated by various biotechnological protocols (ChIP-Seq, RNA-Seq, genome assembly etc.), has grown dramatically and is expected to continue to increase faster than the cost per capacity of storage devices will decrease. This poses challenges for the existing sequence analysis pipelines. They are usually designed to run on a few recent samples, but cannot be used in reasonable time for thousands of samples which makes it very costly to reanalyze existing data. Hence, we are collecting data that grows exponentially in size and are unable to reuse it in its entirety. Searching an entire database for relevant samples enables researches to increase their sample size, where sample size is often a problem, and might even identify new relationships yet unknown. Limiting analysis to recently published datasets is a tremendous waste of resources and a large bottleneck for biomedical research.

The most basic task of such pipelines is to (approximately) search sequencing reads or short sequence patterns like genes in large reference data sets. This has led researchers to develop novel indexing data structures ([SK16] and shortly afterwards [SHCM16],[PAB^+^18], [BdBR^+^19], and [HM20]) to search massive collections of sequences such as RNA-Seq files, pangenomes, or the bacterial and viral metagenome consisting of tens of thousands of species. The most recent tools tackling this problem are Mantis [PAB^+^18], Bifrost [HM20] and Raptor [SMD^+^21].

Mantis relies on *k*-mer counts and uses a counting quotient filter (CQF) to create a space- and time-efficient index for searching queries in multiple samples. The samples are stored in a colored de Bruijn graph, each color (also known as bin) representing a respective sample. Bifrost’s index similarly consists of a colored de Bruijn graph, but uses minimizers and blocked Bloom filters. The Reinert lab introduced a comparable data structure in 2018, the *Interleaved Bloom Filter* (IBF [DSP^+^18a]). Based on this, the application *Raptor* [SMD^+^21] extends the data structure with winnowing minimizers and probabilistic thresholding. Raptor’s performance is currently the state-of-the-art. However, the method has two limitations. Firstly, the runtime deteriorates when the number of samples becomes large. This is the case for metagenomics data, large collections of RNA-Seq files, or indexing *k*-mers in contigs of a de Bruijn graph, among others. Secondly, the size of the index depends on the *maximum* sample size in the collection. As a result, the method uses more space than necessary when the samples are unevenly sized.

In this work we introduce a new data structure, the *Hierarchical Interleaved Bloom Filter* (HIBF) that overcomes the main limitations of the IBF data structure. The HIBF successfully decouples the user input from the internal representation enabling it to handle unbalanced size distributions and millions of samples. This is achieved by computing a hierarchical layout that considers the sequence similarity between bins and automatically sets internal parameters. The resulting data structure can index 98.8 GiB of RefSeq data in less than 13 minutes and is orders of magnitudes faster than comparable tools like Mantis and Bifrost while achieving a comparable or better accuracy. In addition, the index can be built in only 34.4 GiB of space using minimizers, which is 14 times less than the size Mantis uses. The approximate search is up to 129 and 73 times faster than Bifrost and Mantis, respectively. Also, the number of samples that can be used is basically arbitrarily large. We exemplified this by using one million bins, a number is infeasible for previously published tools. This will enable many applications to distribute approximate searches onto very large data sets that occur in metagenomics, pangenomics or sequence archives.

## 2 Results

The data structures and tools mentioned in the introduction explicitly or implicitly address the following problem: Given a set of input sequences (samples), determine in which samples a query sequence can be found with up to a certain number of errors, also known as *Approximate Membership Queries* (AMQ). The most common approach is to store a representation of the samples’ sequence content in an index, which can answer whether a query belongs to one of the input sequences based on sequence similarity. To give an example, consider an input of 25.000 sets of bacterial species, where each set contains the sequences of genomes of all the strains of the respective species. An index over this input can answer whether a query likely originates from one of the 25.000 species. In recent research, each set is called a *color* [PAB^+^18] or *bin* [SMD^+^21]. We will use the term *bin*.

For efficiency, the sequence content is often transformed into a set of representative *k*-mers. The term “representative” indicates that the original *k*-mer content might be transformed by a function that changes its size and distribution. For example, by using winnowing minimizers [MPB^+^17] on the sequence and its reverse complement, or by using gapped *k*-mers ([CLM16]). A winnowing minimizer, which we call (*w, k*)*-minimizer*, is the lexicographically smallest *k*-mer of all *k*-mers and their reverse complements in a window of size *w*. When searching in an index, the same transformation is applied to the corresponding *k*-mers of the query.

### 2.1 The HIBF

The Interleaved Bloom Filter (IBF) data structure published in [SMD^+^21] is a building block in the proposed Hierarchical Interleaved Bloom Filter (HIBF). Its general idea is depicted in Figure 1, further details can be found in [SMD^+^21].

**Figure 1:**
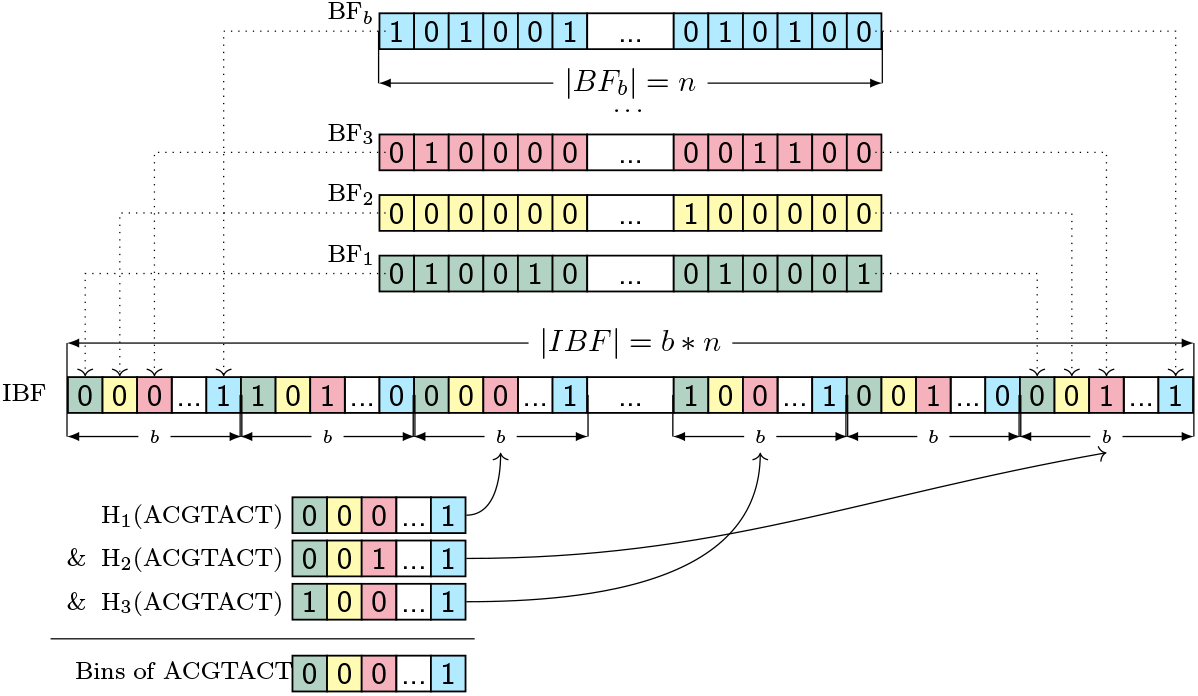
Example of an interleaved Bloom filter (IBF). Differently colored Bloom filters (BF) of length *n* for *b* bins (samples) are shown in the top part. Interleaving the individual Bloom filters yields an IBF of size *b* × *n*. In the example, we use three different hash functions to query a *k*-mer (ACGTACT) and retrieve 3 sub-bitvectors. By combining the sub-bitvectors with a bitwise `&`, we retrieve the *binning bitvector*, where a 1 indicates the presence of the *k*-mer in the respective bin.

The IBF data structure has two limitations. Firstly, due to the interleaved nature of the IBF, the individual Bloom filters must have the same size. Consequently, the largest Bloom filter determines the overall size of the IBF if we want to guarantee a maximal false positive rate (figure 2 a). Consequently, for small-sized bins, the relatively large Bloom filters waste a lot of space. Secondly, although retrieving and combining sub-bitvectors is very efficient in practice, this only holds when the number of bins varies from a few hundreds to a few thousands. An increasing number of user bins will slow down the query speed.

**Figure 2:**
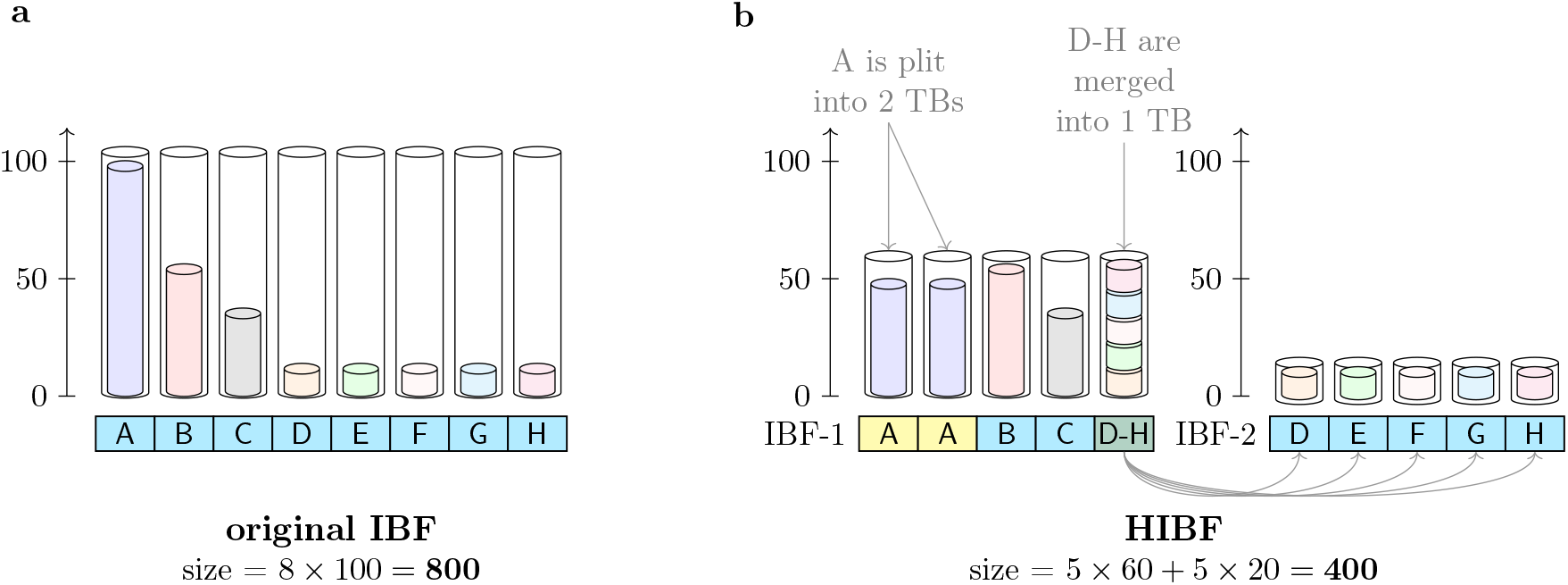
IBF vs. the HIBF. Given an input of eight *user bins* (UBs), subfigure a) displays the layout of a normal IBF storing the content of the UBs, represented by the inner, lightly colored cylinders, in one *technical bin* (TB) each. The outer cylinders represent the size of the TBs and visualizes the wasted space. The horizontal rectangular bar represents the layout, indicating which UBs, identified by their ID (A-H), are stored in which TB. The same semantics hold for subfigure b) which displays the corresponding HIBF with a maximum of 5 TBs on each level. In this example, UB A is split into the first two TBs in IBF 1, while UBs D-H are merged into the last TB. The *merged bin* requires a second lower-level IBF (IBF 2) storing the content of UB D-H in individual TBs. The size is given in exemplary numbers.

The limitations of the IBF arise from the fact that the number of bins and their size directly determines its internal structure. To gain more independence from the input data, the Hierarchical Interleaved Bloom Filter (HIBF) explicitly distinguishes between user bins (UB) and technical bins (TB).

A *user bin* is equivalent to the former term *bin*, namely a set of sequences imbued with a semantic meaning by the user, e.g., all genomes of the species *E. coli* could be in one user bin. Internally, the HIBF stores multiple IBFs on different *levels*. The bins of those IBFs are called *technical bins*. Technical bins may store parts of a user bin or contain the content of multiple user bins. In contrast, in the IBF a user bin is also a technical bin, or simply bin.

The main idea of the HIBF is to *split* large user bins into several technical bins and *merge* small user bins into a single technical bin to even out the *k*-mer content distribution of the technical bins and thereby optimize the space consumption in an IBF (figure 2 b). Specifically, we will either:

1. *split* the *k*-mer content of a user bin and store it in several technical bins,
2. store the entire *k*-mer content of a user bin in a *single* technical bin, or
3. store the *k*-mer content of a range of user bins in one, *merged* technical bin.

Splitting large user bins allows us to lower the maximum technical bin size, while merging small user bins avoids wasting space. However, we cannot use merged bins without further effort because when a query is contained in a *merged bin*, we cannot directly determine in which of the individual user bins the query is. For this purpose, we recursively add a rearranged IBF for each merged bin with the corresponding merged user bins as input. The resulting collection of IBFs and their interconnectivity is what we call the Hierarchical Interleaved Bloom filter, its detailed structure stored in a *layout* (file). To compute a meaningful layout, we engineered a *dynamic programming* (DP) algorithm that optimizes the space consumption of the HIBF data structure (online methods section 4.1.1) given HyperLogLog size estimates of the input data (online methods section 4.1.2). From the layout, we can build the corresponding HIBF index (online methods section 4.1.4), as exemplified in Figure 3. Notably, every IBF in the HIBF except the top-level IBF stores redundant information. In Section 2.2 we show that the space reduction on the top-level (figure 2 b IBF-1) usually compensates the extra space needed for lower levels.

**Figure 3:**
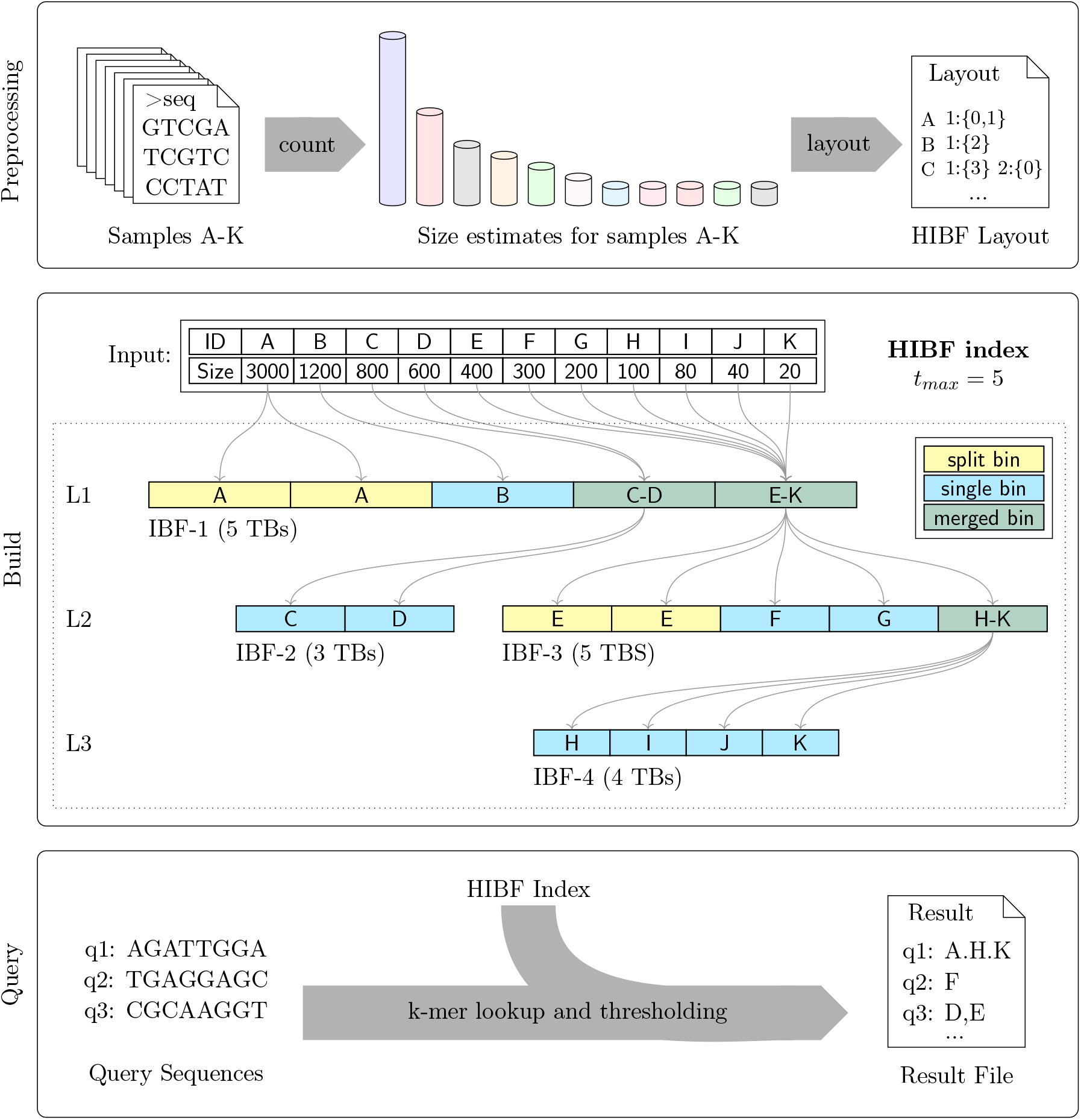
Workflow with details on the HIBF Index structure. Using the HIBF is done in three steps. (1) Preprocessing: Based on the size (representative k-mer content) of the input sequences, a layout is computed with the tool Chopper (section 5). (2) Build: From a given layout, Raptor (section 5) builds an HIBF index depicted in the middle box. (3) Query: The HIBF index can then be used repeatedly to query the membership of sequences inside the input samples. The exemplary HIBF with *t*_*max*_ =5 on 11 *user bins* (UB-A to UB-K) has 3 levels. The first level (L1) is always a single IBF (IBF-1) with exactly *t*_*max*_ *technical bins* (TB). This IBF stores the full data set, structured in a way such that its size is significantly smaller than that of a normal IBF. The individual boxes inside an IBF represent its TBs, which store the *k*-mer content of the labelled *user bin(s)*. For example, in IBF-1 the content of UB-A is stored in two TBs (split), UB-B is stored in one TB (single), and UB-C to UB-D as well as UB-E to UB-K are collected in one TB each (merged). Subsequent levels may have several IBFs. Specifically, they will have one IBF for each *merged bin* on the previous level. For example, on the second level (L2), IBF-2 and IBF-3 correspond to the first and second merged bin of IBF-1, respectively. Note that the IBFs in the layout form a tree, where the root is the top-level IBF and the leaves are formed by IBFs without merged bins.

Since the number of technical bins in each IBF of the HIBF is now independent of the number of user bins, we can choose a relatively small, fixed maximum number of technical bins *t*_*max*_ which allows to efficiently query each IBF. We look up the *k*-mers of a query and apply a refined threshold on each level (online methods section 4.1.5). Traversing lower levels is only required when a query is contained in a *merged bin*. In Section 2.2, we show that, on average, this seldomly happens and that the positive impact of fixing the number of *technical bins* greatly outweighs the disadvantage of searching several levels.

### 2.2 Validation

We incorporated the Hierarchical Interleaved Bloom Filter (HIBF) data structure in the tool *Raptor* [SMD^+^21] that can now be used with either the original IBF or the HIBF. We validated the HIBF data structure on three data sets to show the superiority in comparison to its predecessor (the simple IBF [SMD^+^21]) as well as Mantis and Bifrost: (1) The simulated data set introduced in [SMD^+^21] with more *user bins* (65, 536 versus 64 and 1024 in the original paper), (2) a similarly simulated data set of a million user bins to emphasize the HIBF’s ability to use any number of user bins, and (3) real-world data taken from the RefSeq database [OWB^+^16]. All benchmarks were performed on an Intel Xeon Gold 6248 CPU using 32 threads (supplementary section 7.1).

In order not to give us an advantage, we chose the same parameters proposed by Mantis or Bifrost whenever possible. To this end, the benchmarks were computed with canonical 20-mers (as used by Mantis) and with (24, 20)-minimizers, similar to their usage in Bifrost. In section 2.2.2, we additionally investigated how the HIBF performs with different *k*-mer and minimizer values.

The results of all tools were made comparable in a post-processing step (section 4.2.2) and validated against a ground truth, which, for the RefSeq dataset, was obtained by mapping the queries using a standard read mapper [DSP^+^18b].

#### 2.2.1 Simulated data

Following the approach in [SMD^+^21] we created 64 GiB of data mimicking a metagenomic data set of *b* = 2^16^ = 65, 536 homologous species (*user bins*) and from those ten million queries mimicking a sequencing experiment (supplementary section 7.2.3). This artificial data set is well-balanced, which is the ideal case for the normal IBF. We constructed the (H)IBFs with canonical 20-mers and (24, 20)-minimizers and two different maximal FPRs, 5% and 1.5% (Table 1). For the HIBF *t*_*max*_ was set to 256, the square root of the number of user bins, proven to work well (section 4.3.1). The results show that besides the preprocessing time, the HIBF outperforms Mantis and Bifrost in all cases.

**Table 1:** Benchmark results on simulated data of 65,536 *user bins*. The data compose 64 GiB of random DNA sequence split into 65, 536 *user bins* and ten million queries of length 250 bp containing up to two errors. *Pre-build* includes all steps that have to be done before the actual index is built (online methods section 4.2.1). *Build* measures the time to construct the index, including I/O of all necessary data. *Query* is the multithreaded wall clock time for answering ten million queries. *Statistics* contains the number of false positive and false negative query answers (i.e., whether a read is contained with up to 2 errors).

| Method | Pre-build |  | Build |  |  | Query |  | Statistics |  |
| --- | --- | --- | --- | --- | --- | --- | --- | --- | --- |
|  | Time | RAM | Time | RAM | Index Size | Time | RAM | FP | FN |
| **20-mers** |  |  |  |  |  |  |  |  |  |
| Mantis | 64:10 | 460 MiB | 346:04 | 422 GiB | 29.0 GiB | 35:01 | 29 GiB | 0 | 0 |
| IBF 5% | none |  | 34:58 | **4.1 GiB** | **4.1 GiB** | 24:14 | **7.2 GiB** | 0 | 0 |
| IBF 1.5% | none |  | 42:52 | 8.1 GiB | 8.1 GiB | 18:06 | 11.2 GiB | 0 | 0 |
| 256-HIBF 5% | 103:39 | 829 MiB | **2:31** | 16.8 GiB | 8.1 GiB | **1:18** | 11.6 GiB | 0 | 0 |
| 256-HIBF 1.5% | 100:22 | 827 MiB | 2:42 | 19.5 GiB | 23.5 GiB | 1:20 | 19.5 GiB | 0 | 0 |
| **(24,20)-minimizer** |  |  |  |  |  |  |  |  |  |
| Bifrost | none |  | 143:52 | 9.8 GiB | 4.3 GiB | 54:02 | 75 GiB | 0 | 0 |
| IBF 5% | none |  | 19:46 | **1.6 GiB** | **1.6 GiB** | 7:42 | **4.8 GiB** | 0 | 0 |
| IBF 1.5% | none |  | 17:03 | 3.1 GiB | 3.1 GiB | 5:53 | 6.2 GiB | 0 | 0 |
| 256-HIBF 5% | 100:28 | 829 MiB | **1:18** | 5.6 GiB | 2.7 GiB | **0:46** | 6.0 GiB | 0 | 0 |
| 256-HIBF 1.5% | 104:20 | 827 MiB | 1:25 | 7.4 GiB | 5.1 GiB | **0:46** | 8.5 GiB | 0 | 0 |

Our index based on *k*-mers is between a factor of 1.23 (HIBF 1.5%) to 3.58 times (HIBF 5%) smaller than that of Mantis, which additionally has a very high RAM footprint of more than 400 GiB during building. When using minimizers, the index size can be further decreased to only 2.7 GiB (HIBF 5%) which less than the 4.3 GiB used by Bifrost. For this data set, the HIBF uses more space than the IBF. This is expected, since the data set is ideal in terms of space efficiency for the (original) IBF. While Mantis’s and Bifrost’s index build times are about 6 and 2.5 hours respectively, the HIBF is orders of magnitudes faster, needing only 1 to 3 minutes.

Regarding query runtime, the HIBF is searchable in less than two minutes. For this workload, the best performing HIBF (5%, minimizers) improves on the IBF by a factor of 11. Compared to Mantis and Bifrost, it is faster by a factor of 32 and 71, respectively.

We observed that all tools have no false positive or false negative answers. This is because the randomly simulated DNA sequences of different bins are very dissimilar. Without duplicated *k*-mers, the chance of errors is low.

To emphasize the benefit of the HIBF vs. the IBF for large number of *user bins*, we also constructed a data set with *b* = 2^20^ = 1, 048, 576 *user bins* in the same way (supplementary section 7.2.3). For this large number of *user bins*, the HIBF outperforms the IBF up to a factor of 269 in query time while using only twice as much space. Neither Mantis nor Bifrost could handle this number of *user bins*.

#### 2.2.2 Metagenomic data - RefSeq

This data set consists of 25, 321 files containing the complete genomes of all Archaea and Bacteria in the RefSeq database [OWB^+^16]downloaded using the *genome_updater* (section 6) and has an uncompressed size of about 98.8 GiB. Contrary to the simulated data set, where all *user bins* have the same size, this real-world data set is heavily unbalanced, i.e., the species *Escherichia coli*, represented by 634 assemblies, accounts for almost 7% of all base pairs [PDS^+^20]. From the genomes, we simulated ten million queries. The number of queries per genome is proportional to its size, mimicking sequencing experiments. The benchmarks were computed with canonical 32-mers and (24, 20)-minimizers, as well as two different maximal FPRs (5% and 1.5%) for the (H)IBF. In contrast to section 2.2.1, we chose 32-mers because *Mantis* exited with an error for 20-mers. For the HIBF *t*_*max*_ was set to 192 (section 4.3).

For this data set, the HIBF outperforms all others in index size and query time, with a comparable or better false positive/negative rates (table 2). The HIBF index is built in about 60 minutes (*k*-mers) or 13 minutes (minimizers), taking only slightly longer than the original IBF. Hence the fastest version (HIBF 5%, minimizers) is up to 211 times and 81 times faster than Mantis and Bifrost, respectively, which both need several hours. The factor is much bigger than for the simulated, balanced data set.

**Table 2:**
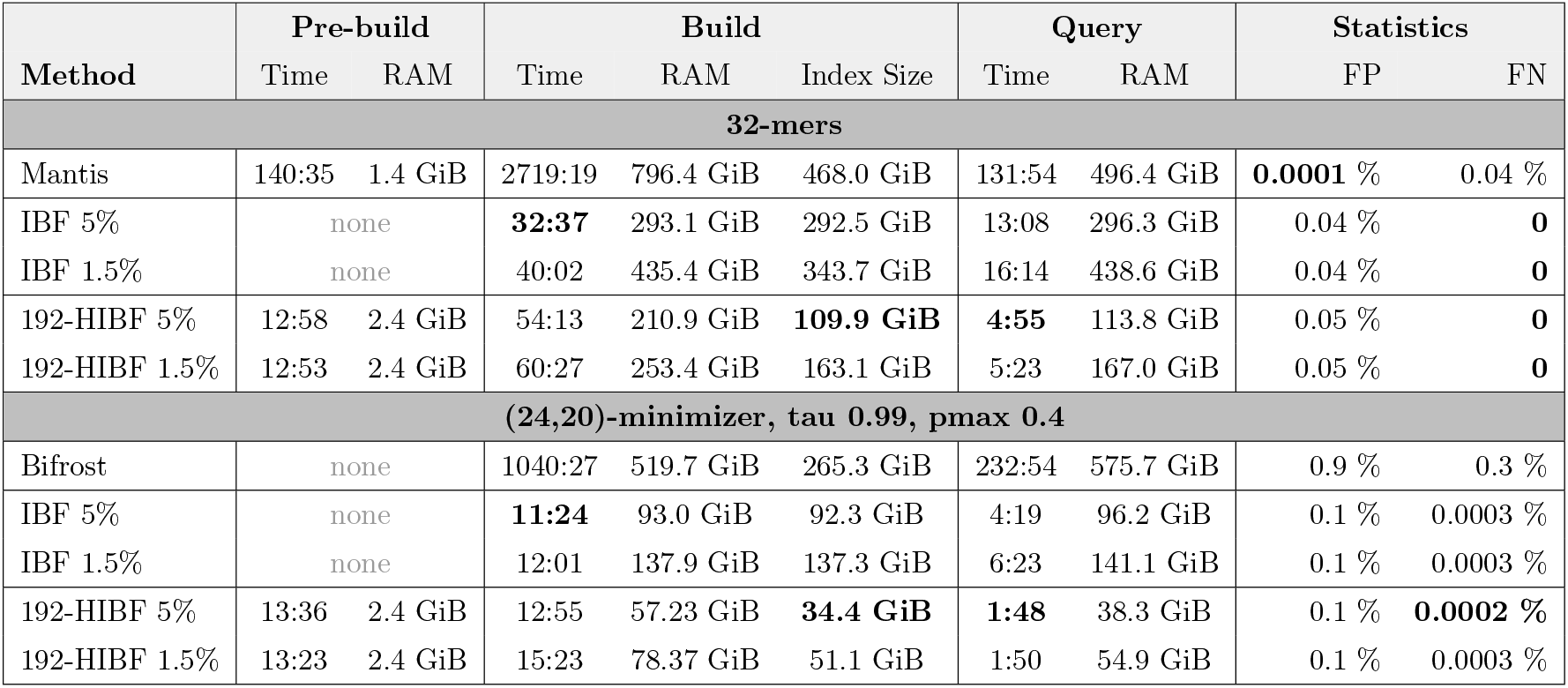
Benchmark results on 25,321 genomes from RefSeq. The uncompressed data set has a size of about 98.8 GiB. Query reads of length 250 *bp* were simulated using the *Mason simulator* [Hol10]. *Pre-build* includes all steps that have to be done before the actual index is built (online methods section 4.2.1). *Build* measures the time to construct the index, including I/O of all necessary data. *Query* is the multithreaded wall clock time for answering ten million queries. *Mantis* does not support minimizers and could only be used with *k* = 32 because it crashed for *k* = 20. Although *Mantis* is technically an exact method outputting *k*-mer counts, a threshold (in this case 0.7) needs to be applied to determine the query membership resulting in few false positives/negatives. Bifrost was run with (24, 20)-minimizers and a threshold of 0.36. For details on the chosen thresholds, see section 2.2.

| Method | Pre-build |  | Build |  |  | Query |  | Statistics |  |
| --- | --- | --- | --- | --- | --- | --- | --- | --- | --- |
|  | Time | RAM | Time | RAM | Index Size | Time | RAM | FP | FN |
| **32-mers** |  |  |  |  |  |  |  |  |  |
| Mantis | 140:35 | 1.4 GiB | 2719:19 | 796.4 GiB | 468.0 GiB | 131:54 | 496.4 GiB | **0.0001** % | 0.04 % |
| IBF 5% | none |  | **32:37** | 293.1 GiB | 292.5 GiB | 13:08 | 296.3 GiB | 0.04 % | **0** |
| IBF 1.5% | none |  | 40:02 | 435.4 GiB | 343.7 GiB | 16:14 | 438.6 GiB | 0.04 % | **0** |
| 192-HIBF 5% | 12:58 | 2.4 GiB | 54:13 | 210.9 GiB | **109.9 GiB** | **4:55** | 113.8 GiB | 0.05 % | **0** |
| 192-HIBF 1.5% | 12:53 | 2.4 GiB | 60:27 | 253.4 GiB | 163.1 GiB | 5:23 | 167.0 GiB | 0.05 % | **0** |
| **(24,20)-minimizer, tau 0.99, pmax 0.4** |  |  |  |  |  |  |  |  |  |
| Bifrost | none |  | 1040:27 | 519.7 GiB | 265.3 GiB | 232:54 | 575.7 GiB | 0.9 % | 0.3 % |
| IBF 5% | none |  | **11:24** | 93.0 GiB | 92.3 GiB | 4:19 | 96.2 GiB | 0.1 % | 0.0003 % |
| IBF 1.5% | none |  | 12:01 | 137.9 GiB | 137.3 GiB | 6:23 | 141.1 GiB | 0.1 % | 0.0003 % |
| 192-HIBF 5% | 13:36 | 2.4 GiB | 12:55 | 57.23 GiB | **34.4 GiB** | **1:48** | 38.3 GiB | 0.1 % | **0.0002** % |
| 192-HIBF 1.5% | 13:23 | 2.4 GiB | 15:23 | 78.37 GiB | 51.1 GiB | 1:50 | 54.9 GiB | 0.1 % | 0.0003 % |

Comparing the resulting index sizes, the advantage of the HIBF becomes even more pronounced. Using 32-mers, Mantis has an index size of almost 500 GiB and the original IBF about 300 GiB (5%) or 350 GiB (1.5%) whereas the HIBF only needs a little more than 100 GiB (5%) or less than 200 GiB (1.5%). The HIBF index is almost as small as the original data set. When using minimizers, the index size can be further reduced to 34 GiB, only a fraction of the original input. This is a third of the original IBFs size and an 8-fold improvement compared to Bifrost. As expected, a lower maximum false positive rate results in higher index sizes for the (H)IBFs.

Most interestingly, regarding query times, the fastest version of the HIBF (HIBF 5%, minimizers) surpasses Mantis by a factor of 73 and Bifrost by a factor of 129. The HIBF needs between 2 (*k*-mers) and 6 (minimizers) minutes for the 10 million queries, whereas Mantis needs about two hours and Bifrost almost four hours. It improves on the original IBF by a factor of 3. Additionally, the output of Mantis encompasses about 400 GiB, that of Bifrost 473 GiB, whereas Raptor using the (H)IBF only produces a file of 5 GiB in size (supplementary section 7.2.1).

If no minimizers are used, our thresholding ensures that the (H)IBF gives no false negative answers. In contrast, Mantis has a false negative rate of 0.04%. Using minimizers, there are few false negatives, because the minimizer compression is not lossless. The false positive rates are higher than that of Mantis, but usually tools can easily handle false positive by a subsequent validation step, while false negatives can be troublesome.

We additionally investigated the influence of varying values for *k* and (*w, k*) for the HIBF (1.5%) in terms of index size, query time and accuracy (figure 4). For comparison, we included the results of table 2 for Mantis, Bifrost, and the IBF 1.5%.

**Figure 4:**
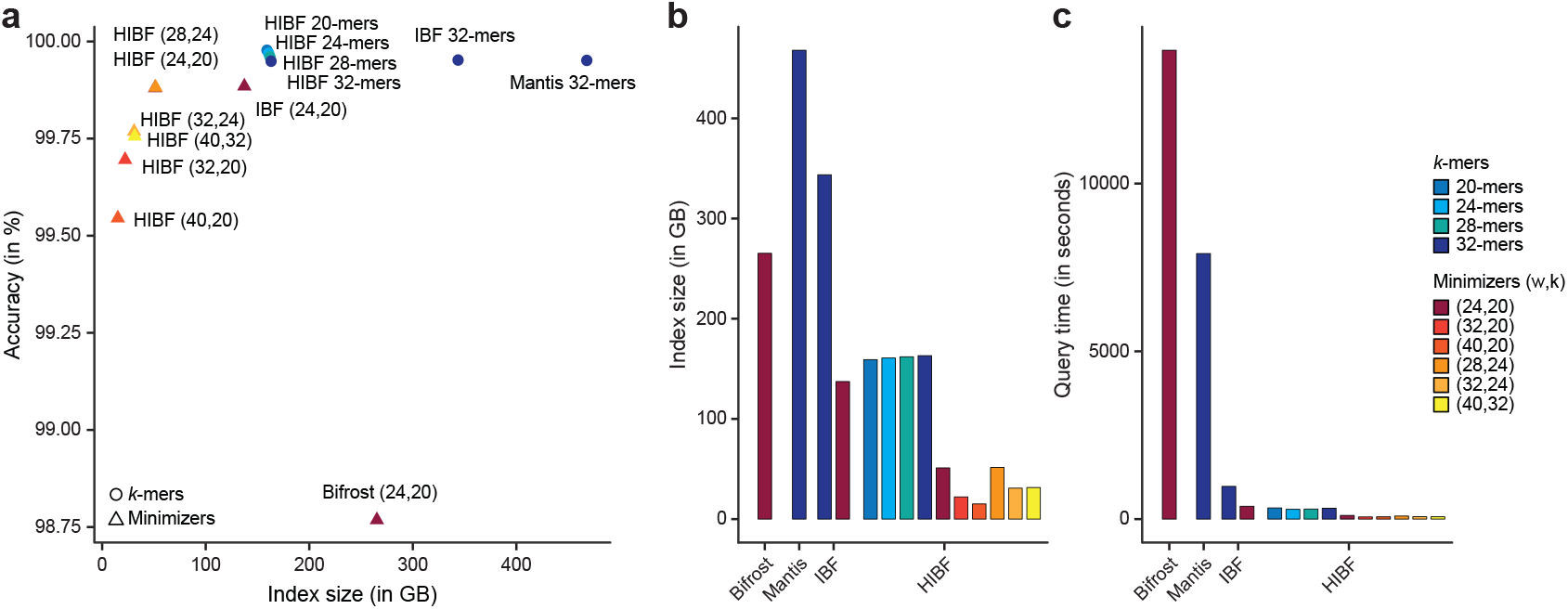
Additional experiments with varying k and (w,k) for the HIBF. The maximum false positive rate for the (H)IBF was fixed to 1.5% and *t*_*max*_ = 192 for the HIBF. **a** For an HIBF using canonical *k*-mers, the choice of *k* had small impact on index size and accuracy and achieved the best overall accuracy. Compressing the index with minimizers decreased the accuracy. Although the accuracy is, in general, excellent, with > 98% for the least performing tool Bifrost, it has to be noted that it is biased by a high number of true negatives and thus small differences in accuracy have a high impact on the actual numbers of false positives/negatives. **b** The index size in GiB and **c** the query time in seconds of the experiments on the RefSeq data set.

Figure 4 shows that choosing different values for *k* when using canonical *k*-mers did not significantly affect the index size, query time nor accuracy of the HIBF. The best accuracy was achieved with 20-mers which is marginally better than Mantis but uses only a third of its index size. Using minimizers could largely reduce the index size and query time, but had a negative impact on the accuracy. The larger the difference of *w* to *k*, the higher is the compression, which lowers the threshold for the query membership. This resulted in an increasing number of false positives, thus decreasing the accuracy. False negatives were rare and only slightly affected by the choice of (*w, k*) (supplementary table 6).

## 3 Discussion

In this work, we presented a general data structure and method to store a representative set of *k*-mers of a data set that is partitioned into hundreds up to millions of user bins in such a way, that very fast approximate membership queries are possible. The HIBF data structure has enormous potential. It can be used on its own like in the tool Raptor, or can serve as a prefilter to distribute more advanced analyses such as read mapping.

Since the build time exceeds two orders of magnitude less than that of comparable tools like Mantis [PAB^+^18] and Bifrost [HM20], the HIBF can easily be rebuilt even for huge data sets. In our experiments, we could index 98.8 GiB of RefSeq data (Archaea and Bacteria complete genomes as of 28-01-2022) in less than 13 minutes using 34.4 GiB of space. This is 211 times faster than Mantis, while at the same time we use 14 times less space. The approximate search is up to 73 times faster than Mantis and 129. times faster than Bifrost.

Moreover, the number of user bins (resp. colors) that can be used is almost arbitrarily large. We exemplified this by using one million bins, a number not possible for any other published tool. In this extreme scenario, it took just under a day to calculate the layout. Still, we can foresee future improvements in this algorithm to speed up the layout computation and allow even more bins to be used in reasonable time.

If we assume that a HIBF index needs about 40% of the input size, and that we can handle millions of user bins, one can use the HIBF to index very large sequence repositories. For example, the European Nucleotide Archive (ENA) contains currently about 2.5 million assembled/annotated sequences with a total of ≈ 11 Tbp [CAA^+^22]. Using the HIBF data structure, we could build 11 HIBF indices, each storing roughly 1 TB of sequence content in about 200, 000 *user bins*. We would expect each index to have a size of about 400 GiB which easily fits into the main memory of a server. We also showed that hundreds of thousands of *user bins* can be readily handled by the HIBF. Subsequently, querying ten million reads could be done by querying 11 HIBFs on different machines in parallel in less than 2 minutes. A handful of queries would be answered in a fraction of a second. Hence, in this setting the archives could offer a portal for individual user queries, knowing that they can answer about 100, 000 queries (of length 250) per second.

## 4 Online Methods

The following sections give the details referred to in the Results section. We will use the following notation:

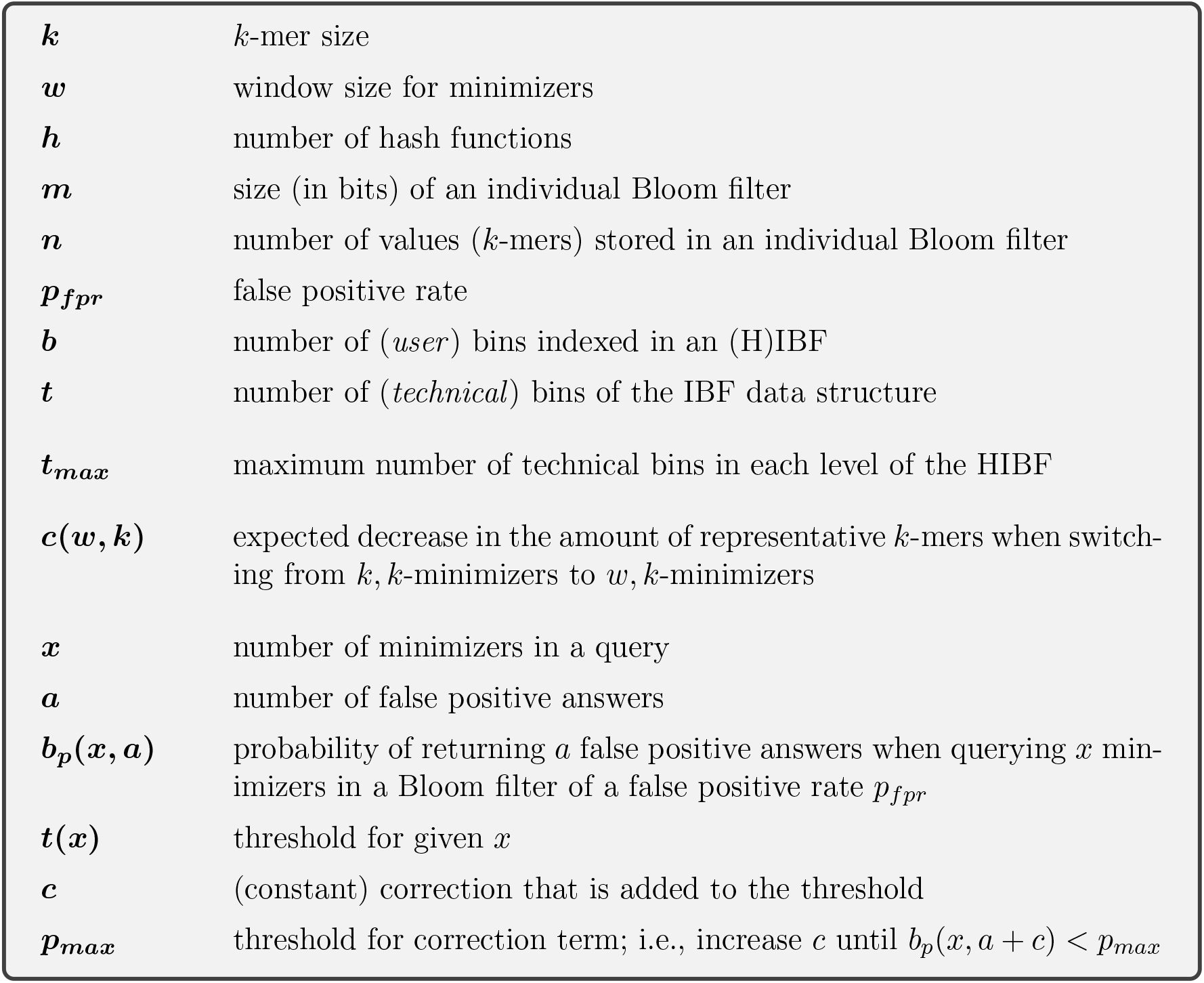

|  |  |
| --- | --- |
| $k$ | $k$-mer size |
| $w$ | window size for minimizers |
| $h$ | number of hash functions |
| $m$ | size (in bits) of an individual Bloom filter |
| $n$ | number of values ($k$-mers) stored in an individual Bloom filter |
| $p_{fpr}$ | false positive rate |
| $b$ | number of (*user*) bins indexed in an (H)IBF |
| $t$ | number of (*technical*) bins of the IBF data structure |
| $t_{max}$ | maximum number of technical bins in each level of the HIBF |
| $c(w, k)$ | expected decrease in the amount of representative $k$-mers when switching from $k, k$-minimizers to $w, k$-minimizers |
| $x$ | number of minimizers in a query |
| $a$ | number of false positive answers |
| $b_p(x, a)$ | probability of returning $a$ false positive answers when querying $x$ minimizers in a Bloom filter of a false positive rate $p_{fpr}$ |
| $t(x)$ | threshold for given $x$ |
| $c$ | (constant) correction that is added to the threshold |
| $p_{max}$ | threshold for correction term; i.e., increase $c$ until $b_p(x, a + c) < p_{max}$ |

### 4.1 The HIBF

#### 4.1.1 Computing an HIBF Layout - a DP algorithm

In this section, we explain how to compute a layout, i.e., how we decide which user bins to split and which to merge (Figure 3). To this end, we engineered a dynamic programming (DP) algorithm that heuristically optimizes the space consumption of a single IBF, considering the space needed for lower level IBFs for *merged bins*. Thus, computing the layout on the first, main level IBF optimizes its layout while estimating the size of the entire HIBF.

As outlined in section 2.1, we then recursively apply the DP algorithm on each lower level IBFs.

Assume that we have *b* user bins UB_0_,…, UB_*b*−1_ sorted in decreasing order of their size, |UB_0_|≥ … ≥|UB_*b*−1_|, which we want to distribute to *t* technical bins TB_0_,…, TB_*t*−1_ in a single IBF. While not strictly necessary, the *order* is chosen such that small user bins cluster next to each other because in our algorithm only contiguous bins may be merged. We denote the *union* size estimate of a *merged bin* that stores the *k*-mer multiset UB_*i*_ ∪ UB_*i*+1_ … ∪ UB_*j*_ as *U*_*i,j*_. We use HyperLogLog sketches [FF07] (for details, see 4.1.2) to quickly estimate the size of the union when merging user bins. Since the IBF only stores a representative *k*-mer’s presence (unique *k*-mer content), not how often it was inserted, the size of the *merged bin* may be smaller than the sum of sizes of the respective user bins. The effect is that merging user bins of similar sequence content results in smaller sized *merged bins* which is beneficial for the overall size of the IBF. We exploit this advantage by the optional step of rearranging the user bins beforehand based on their sequence similarity (see section 4.1.2).

Further, the user needs to fix the number of hash functions *h*, the desired false positive rate *p*_*fpr*_, and the maximum number *t*_*max*_ of technical bins on each IBF of the HIBF. The former two parameters are needed to estimate the IBF size and must correspond to the parameters of building the index afterwards. The latter is critical for the HIBF layout. Section 4.3.1 discusses sensible defaults.

Regarding the expected false positive rate of the HIBF index, we point out that when splitting a user bin we introduce a *multiple testing problem*. This happens because we query a *k*-mer for a split bin several times in several technical bins. We correct for this by increasing the size of the respective technical bins by a factor *f*_*corr*_(*s, p*_*fpr*_) where *s* is the number of technical bins the user bin is split into and *p*_*fpr*_ is the desired false positive rate (see details in 4.1.3).

The *general sketch* of the algorithm is the following: The dynamic programming (DP) matrix *M* has *t* rows, representing TB_0_,…, TB_*t*−1_ and *b* columns, representing UB_0_,…, UB_*b*−1_. When we move horizontally in the matrix, we consume multiple user bins while remaining in a single technical bin. This indicates a merged bin. When we move vertically, we consume multiple technical bins while remaining at a single user bin. This indicates a split bin. We treat a *single bin*, introduced for clarity in section 2.1, as a split bin of size 1. We do not move along the diagonal. The above semantics allow us to verbalize a structure that is then used to estimate the space consumption and compute a local optimum in each cell of the DP matrix. Specifically, the space consumption is tracked using two *t* × *b* matrices *M*_*i,j*_ and *L*_*i,j*_:

- *M*_*i,j*_ tracks the maximum technical bin size 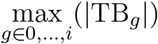 when the first *j* +1 user bins are distributed to *i* +1 technical bins. Minimizing the maximum technical bin size optimizes the IBF space consumption, since it directly correlates with the total IBF size (see section 2.1 and Figure 2).
- *L*_*i,j*_ tracks the space consumption estimate of all lower level IBFs that need to be created for each merged bin given the structure imposed by *M*_*i,j*_.

##### Initialization

For the first column, when *j* = 0, there is only a single user bin UB_0_ which can be split into *t* technical bins. Therefore, the maximum technical bin size stored in *M*_*i*,0_ is the size of UB_0_ divided by the current number of technical bins *i* it is split into, corrected by *f*_*corr*_ (for details on *f*_*corr*_ see section 4.1.3). Since no merging is done, no lower level IBFs are needed and *L*_*i*,0_ is always 0.

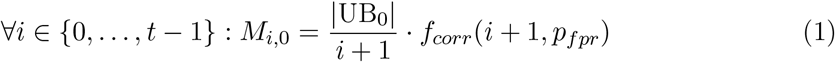

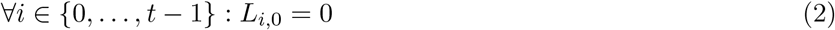

For the first row, when *i* =0 and *j* ≠ 0, all user bins have to be merged into a single technical bin. The size of the resulting *merged bin* is estimated using the precomputed unions *U*_0,*j*_ (see 4.1.2 for details). Since we create a *merged bin* in each step, the resulting additional space consumption on lower level IBFs is estimated by the sum of sizes of contained user bins times the maximal number of levels 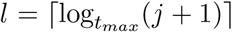. We use the sum instead of the union here because we do not know beforehand how the lower level IBFs are laid out, so we estimate the worst case of storing every *k*-mer again on each lower level.

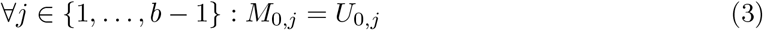

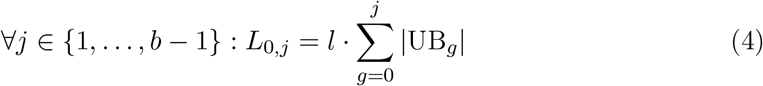

##### Recursion

We store the maximum technical bin size in *M*_*i,j*_ and the lower level costs in *L*_*i,j*_ and we want to optimize the total HIBF space consumption. It is computed by *M*_*i,j*_ times the number of technical bins, which is the size of the first, main level IBF, plus the lower level costs *L*_*i,j*_:

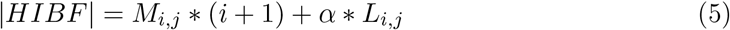

The *parameter α* can be used to tweak the influence of lower levels on the space and query time of the HIBF. The query time decreases when many lower levels are introduced since we have to traverse and query a lot of IBFs, but the space consumption is often lower. When *α* = 1, the space consumption is optimized by expecting our lower level space estimate to be exact. If we choose higher values for *α*, we artificially increase the costs of *merged bins* and their lower levels, thereby decreasing their occurrence in the layout. A sensible default is that was experimentally derived to work well is *α* = 1.2.

In the recursion, we process each cell *M*_*i,j*_, *i* ≠ 0 and *j* ≠ 0, by deciding for the next user bin UB_*j*_ whether to (1) split it into *i* − *i*′, for some *i*′ < *i*, technical bins or (2) merge it with all user bins starting at *j*′, for some *j*′ < *j*.

In the first case, when *splitting UB*_*j*_ thereby moving vertically, we want to find the *i*′ < *i* that results in the smallest overall HIBF size *v*_*i,j*_ (figure 5 c). Semantically, we come from the layout *M*_*i*′,*j* −1_ which already distributed UB_0_,…, UB_*j*−1_ into TB_0_,…, TB_*i*′_, leaving the technical bins TB_*i*′+1_,…, TB_*i*_ to store the split content of UB_*j*_. The new HIBF size is computed analogous to equation 5 with the new maximal technical bin size *m*_*v*_ and the new lower level costs *l*_*v*_. Since we introduce no *merged bin, l*_*v*_ is simply the former costs *L*_*i*′,*j*−1_. *m*_*v*_ is computed by taking the maximum of the former value *M*_*i*′,*j*−1_ and the *split bin* size we obtain from equally dividing the *k*-mer content of UB_*j*_ into *i* − *i*′ technical bins corrected by *f*_*corr*_.

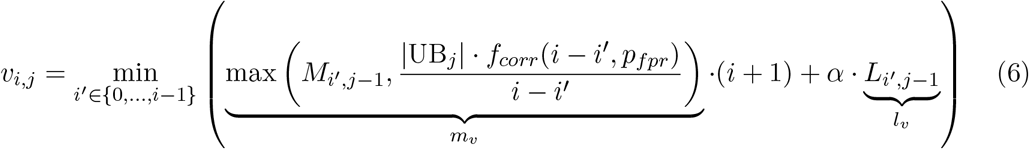

In the second case, when *merging UB*_*j*_ thereby moving horizontally, we want to find the *j*′ < *j* − 1 that results in the smallest overall HIBF size *h*_*i,j*_ (figure 5 b). Semantically, we come from the layout *M*_*i*−1,*j*′_ which already distributed UB_0_,…, UB_*j*′_ into TB_0_,…, TB_*i*−1_, leaving technical bin TB_*i*_ to store the merged content of UB_*j*′+1_,…, UB_*j*_. The new HIBF size is computed analogous to equation 5 with the new maximal technical bin size *m*_*h*_ and the new lower level costs *l*_*h*_. *m*_*h*_ is computed by taking the maximum of the former value *M*_*i*−1,*j*′_ and the *merged bin* size we obtain from the precomputed Union *U*_*j*′+1,*j*_. Since we introduce a new *merged bin l*_*h*_ is computed by adding to the former costs *L*_*i*−1,*j*′_, the space estimate of *l* expected lower levels times the sum of contained user bin sizes (for reasoning see that of equation 4).

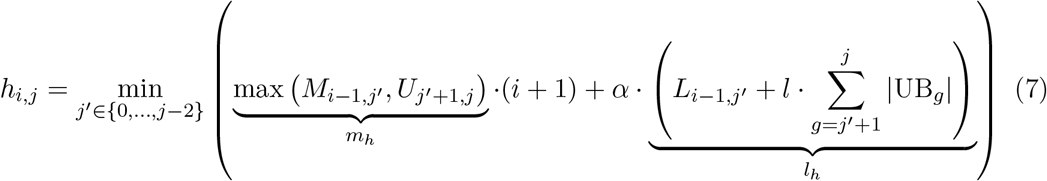

Given *m*_*v*_, *l*_*v*_, *m*_*h*_ and *l*_*h*_ for which the minima in *v*_*i,j*_ and *h*_*i,j*_ are achieved, respectively, the values of the cells *M*_*i,j*_ and *L*_*i,j*_ are updated according to the minimum of *v*_*i,j*_ and *h*_*i,j*_:

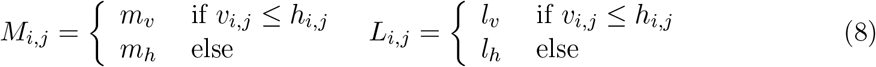

The above recurrence can be used to fill the two dynamic programming matrices row- or column-wise, because the value of a cell depends entirely on cells with strictly smaller indices. The traceback can then be started from the cell *M*_*t*−1,*b*−1_.

**Figure 5:**
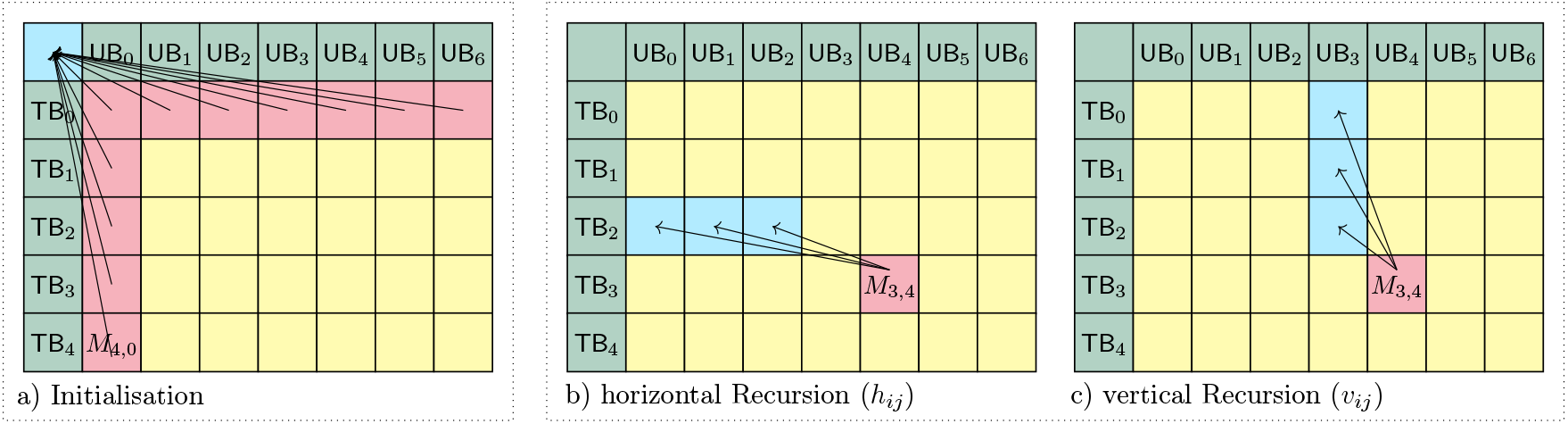
DP Algorithm traceback visualization. Matrices a, b, and c visualize the algorithm for a layout distributing *b* = 7 user bins (columns) to *t* = 5 technical bins (rows). (a) Shows the traceback of the initialization, all coming from the virtual starting point (−1, −1). E.g., *M*_4,0_ represents the sub-layout of splitting UB_0_ into all available technical bins. (b) shows which cells *j*′ < *j* − 1 = 3 are considered in the horizontal Recursion, e.g., *j*′ =2 would indicate to merge UB_3_ and UB_4_ into TB_3_. (c) shows which cells *i*′ < *i* =3 are considered in the vertical Recursion, e.g., *i*′ =1 would indicate to split UB_4_ into TB_2_ and TB_3_.

Since we use HyperLogLog sketches, the algorithms runs quite fast in practice. For example, it takes only 13 minutes to compute the layout with *t*_*max*_ = 192 of user bins for a data set consisting of all complete genomes of archaea and bacteria species in the NCBI RefSeq database [OWB^+^15] (about 25, 000 genomes with total size of roughly 100 GiB).

#### 4.1.2 HyperLogLog estimates

##### HyperLogLog

The *HyperLogLog* (HLL) algorithm [FF07] approximates the number of distinct elements in the input data that were previously converted into a set of uniformly distributed, fixed-size hash values. It is based on the observation that a hash value of length *q* > *p* has a probability of 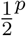 of containing *p* leading zero. When keeping track of the maximum number of leading zeros *p*_*max*_ of the data’s hash values, one can estimate its size by 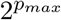. Since this estimate has a very high variance, it is improved by splitting the data into *m* = 2^*b*^, *b* ∈ [4,*q* − 1], subsets and keeping track of *p*_*max*_ for each subset. These *m* values are called a *sketch* of the input data. The size estimate of the original input is computed by using the harmonic mean and bias correction of the *m* values of the subsets (see [FF07] for details).

In our application we convert the input sequences into 64-bit *k*-mers by arithmetic coding, hash them using the third-party function `XXH3_64bits`^1^ and then apply the HLL algorithm with *q* = 64 and *b* = 12 using our adjusted implementation of ^2^.

##### Union estimation

Given two *HyperLogLog* sketches, e.g., created from two user bins, with the same number of values *m*, they can be merged by inspecting the *m* values of both sketches and simply storing the larger (*p*_*max*_) value of each. The resulting new sketch can estimate the size of the combined sketches, namely the number of distinct elements in the union of the two user bins. When merging several *sketches*, we merge the first two and then merge the rest iteratively into the new union sketch.

In our application, when iterating column-wise over the matrix of the DP algorithm, for each column *j* we precompute the *j* − 1 union estimates *U*_*j*′,*j*_ for *j*′ ∈ [0, …, *j* − 1] with

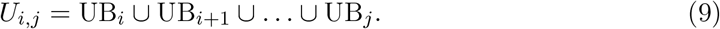

##### Rearrangement of user bins

When merging two or more user bins, we unite their *k*-mer content by storing all unique *k*-mers present in each user bin. It follows that the size of the union is always smaller or equal to the sum of sizes of the united user bins. For a merged bin in the HIBF, this means that we can potentially save space my uniting user bins of similar *k*-mer content. To exploit this advantage, the user has the option to partially rearrange the sorted order of user bins based on their similarity.

We estimate the similarity of two user bins by computing the *Jaccard distance* based on *HyperLogLog* sketches. The *Jaccard distance d*_*J*_ (*A, B*) is defined as 1 minus the *Jaccard index J*(*A, B*) and can be rewritten in the following way:

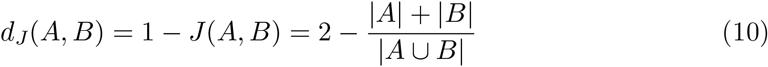

We use the rightmost term to compute the *Jaccard distance* using the union sketch of A and B to estimate | *A*∪*B*|.

When rearranging, we still want to partly maintain the initial order of user bin sizes for two reasons: 1) for the HIBF it is beneficial to merge small user bins (see Figure 2) and 2) a limitation of the DP algorithm is that only user bins next to each other can be merged so ordering them by size clusters small bins together. We therefore only rearrange user bins in non-overlapping *intervals*. The size of an interval is determined by the parameter *r*_*m*_, the maximum ratio of the largest to the smallest user bin within the interval.

Within an interval, we perform an agglomerative clustering on the range of user bins based on their *Jaccard distances*. The order of user bins is switched according to the resulting cluster tree, leaving similar user bins adjacent to each other (see Figure 6).

**Figure 6:**
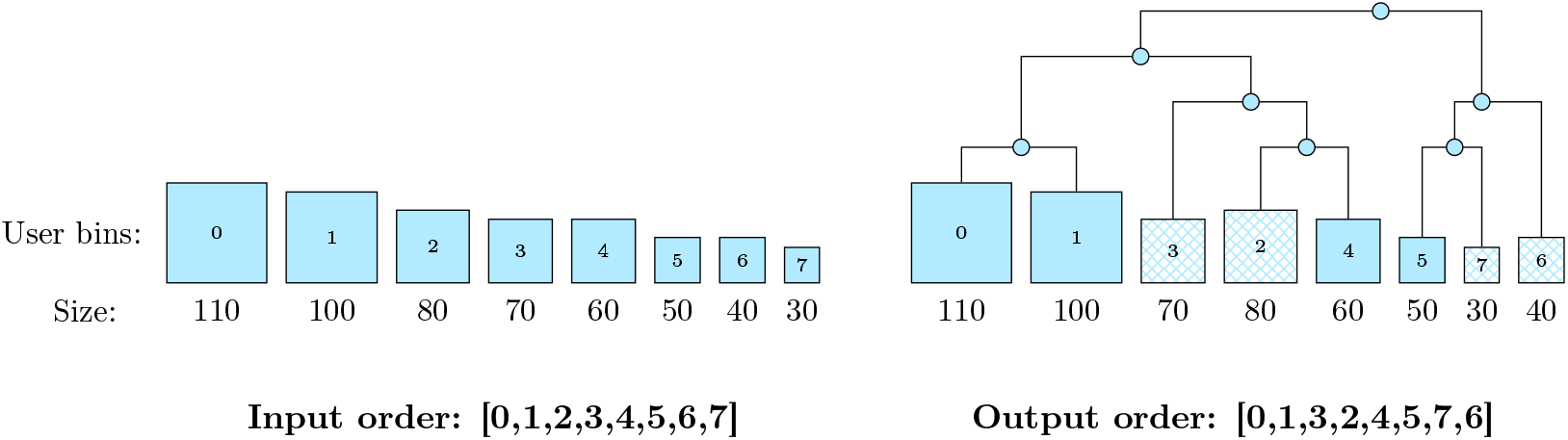
Rearranging user bins by agglomerative clustering. Given eight user bins (UBs) as input, sorted by their size (left-hand side) the example shows how the order may change after applying an agglomerative clustering algorithm based on sequence similarity (right-hand side). The agglomerative clustering arranged UB-2 and UB-4 next to each other, as well as UB-5 and UB-7.

#### 4.1.3 IBF size correction for split bins

Recall that for a given false positive rate *p*_*fpr*_, the size *m* (bits) of an individual Bloom Filter is proportional to the amount of *k*-mers *n* it is designed to store. If *h* is the number of hash functions, then it holds:

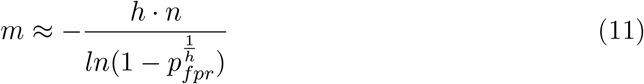

If we split a user bin into *s* technical bins and divide the *k*-mer content equally among them, we cannot simply use the above equation 11 on each to determine its size because we introduce a multiple testing problem. Namely, the probability to obtain a false positive answer for the user bin when querying those *s* technical bins is 1 − (1 − *p*_*fpr*_)^*s*^, since we only get no false positive answer, if *all split bins* have no false positive answer. We can correct the individual false positive rates to reflect the desired overall *p*_*fpr*_, by increasing the size of the *split bins* by the factor *f*_*corr*_(*s, p*_*fpr*_) which is

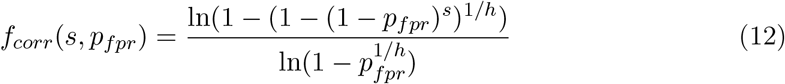

For example, given *h* = 4 and *p*_*fpr*_ = 0.01, if we split a user bin into *s* = 5 technical bins their size computed with equation 11 must be increased by the factor 1.673. If we split the same user bin into *s* = 20 technical bins the size must be corrected by 2.786.

#### 4.1.4 Building an HIBF index from a layout

The hierarchy in an HIBF layout forms a tree (Figure 3). To avoid reading input files several times, we build the index using a *bottom-up* strategy implemented by a recursive construction algorithm. The recursion anchor lies at IBFs without merged bins (leaves of the tree) and traverses the layout similar to a *breadth-first search*. At a leaf, we read the respective input samples and transform them into their representative *k*-mer content.

Based on the *k*-mer contents, we can construct an IBF whose size fits the desired maximum false positive rate. We then insert the *k*-mer contents into the IBF. Moving along the tree, we keep the *k*-mer contents of child nodes to insert them into the respective merged bins of parent IBFs. At a parent node, the input samples from split and single bins are read, processed and, together with the merged bin content from child nodes, used to construct the current IBF. The algorithm ends at the root IBF on level L1. We trivially parallelized the construction of independent lower levels.

#### 4.1.5 Querying an HIBF

To query the HIBF for a sequence, we employ a *top-down* strategy. First, the query sequence is transformed into relevant *k*-mers in the same way as it was done for the index construction. For each *k*-mer, we determine its membership in the first, main level IBF of the HIBF by counting the total number of *k*-mers that occur in each *technical bin*. Subsequently, we gather all *technical bins* whose count is equal to or exceeds a certain *threshold*. For *split bins*, the *k*-mer counts of all bins associated with the respective *user bin* need to be accumulated before thresholding. For *split* and *single bins*, we can directly obtain the corresponding *user bins* to answer the query. For *merged bins* that exceed the threshold, we need to apply the same procedure recursively on the associated child IBF on a lower level. Notably, having a threshold allows us to skip traversing lower level IBFs whose upper level *merged bins* already do not exceed the threshold. In practice, this means that we only have to access a fraction of the HIBF. The final result is the set of all *user bins* that contain a query.

The threshold can be either user-defined or computed for a given amount of allowed errors *e* in the query. For the latter, we rely on the method introduced in [SMD^+^21]. Briefly, we utilize a well-known *k*-mer lemma [JU91]. However, when winnowing minimizers are used, we apply a probabilistic model that accounts for how errors might destroy relevant *k*-mers [SMD^+^21]. The model returns a threshold value given how many relevant *k*-mers are in the query. In this work, we further refined the model to incorporate the effect of the false positive rate on the threshold (online methods 4.3.3). The thresholding step is a convenience for the user missing in tools like Mantis [PAB^+^18].

### 4.2 Validation

#### 4.2.1 Pre-build tasks

The preprocessing encompasses all steps that have to be done before the actual index is built. For *Raptor* with an HIBF, this includes estimating the size of *user bins* using HLL sketches 4.1.2 and computing the layout 4.1.1.

For Mantis, this involves converting the input to FASTQ format (required), running Squeakr [PBJP18], and converting the query into the proprietary format required by Mantis. Bifrost and Raptor using the IBF have no preprocessing steps.

#### 4.2.2 Post-processing results of other tools

The results from Mantis could not be directly compared to that of Raptor because Mantis is missing the thresholding step to finally determine the membership of a query. Mantis solely outputs counts obtained for the unique *k*-mers of each query. We therefore applied a threshold that is in accordance with the *k*-mer lemma [JU91]. For reads of length 250 with up to 2 errors, as in our experiments, this corresponds to a threshold of 0.7 or 155 *k*-mers that must be found. For Bifrost, we chose a threshold of 0.36, calculated using the *k*-mer lemma for *k* = 24, which is the expected average number of minimizers in this setting.

### 4.3 Key parameters

The HIBF has many parameters that influence the memory consumption and run time, and partially depend on each other. We acknowledge that this can be challenging for the user. To alleviate the problem, we will in the following sections (1) describe the impact of the key parameters on the performance of the HIBF and finally (2) give sensible defaults and describe how the HIBF can set them automatically given very few, easy to understand parameters.

#### 4.3.1 The choice of *t*_*max*_

The choice of *t*_*max*_, i.e., the maximum number of technical bins of the individual IBFs in an HIBF, influences the depth of the HIBF which impacts both total index size and query runtime. We show in the followinpg that the optimal *t*_*max*_ is near the square root of the number of user bins *b*, i.e. 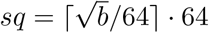

If *t*_*max*_ *is relatively small*, the depth of the HIBF is large, but the individual IBFs will be small. In terms of query runtime, querying a single, small IBF is fast, but we potentially have to traverse many levels. In terms of space, a small *t*_*max*_ and thus a large HIBF depth results in storing a lot of redundant information and thus the space consumption increases. If we choose *t*_*max*_ *relatively large*, the depth of HIBF levels and the number of IBFs will be low, but the individual IBF sizes will be large. In terms of query runtime, we then expect few queries on each level, but querying a single IBF is more costly. In terms of space, the larger *t*_*max*_, the more technical bins we have to cleverly distribute the content of the user bins to and thus lower the space consumption. For some values of *t*_*max*_, the space consumption increases, although *t*_*max*_ is large. This happens because IBFs on the last level contain only a few UBs, which will lead to relatively high correction factors *f*_*h*_ (see Section 4.1.3).

With a simple experiment, we validated one of the above observations, namely that the query runtime of a single IBF increases with the number of (technical) bins (see Table 3). The results show an increase in query runtime for each false positive rate. Additionally, we observe that the higher the false positive rate, the greater the query runtime penalty with increasing number of bins. We can use those results to optimize the choice of *t*_*max*_, which is explained in the following.

**Table 3:** Runtime penalty for increasing number of bins in an IBF. The values are given as ratios by which a single query is more costly when using an IBF with *b* bins compared to an IBF with the minimum number of 64 bins. We simulated *b* equally sized (*user*) bins for *b* ∈ {64, 256, 512, &, 32768} of random sequence content (4.3*GiB* in total) and sample 10 Million reads of length 250. We then constructed a normal IBF over these *b* bins using (24, 20)-minimizers and 4 hash functions. We conducted this experiment three times, for the false positive rates 0.0001, 0.0125 and 0.3125 which resulted in IBFs of different densities. Next, we measured the time required to count the *k*-mer occurrences in each of the *b* bins for all 10 million reads (5 repetitions).

| **b** | **64** | **128** | **256** | **512** | **1024** | **2048** | **4096** | **8192** | **16384** | **32768** |
| --- | --- | --- | --- | --- | --- | --- | --- | --- | --- | --- |
| $p_{fpr} = 0.0001$ | 1.00 | 1.02 | 1.25 | 1.56 | 1.91 | 2.67 | 4.07 | 6.14 | 9.84 | 17.74 |
| $p_{fpr} = 0.0125$ | 1.00 | 1.05 | 1.29 | 1.67 | 2.37 | 3.34 | 5.80 | 9.63 | 17.59 | 33.31 |
| $p_{fpr} = 0.3125$ | 1.00 | 1.15 | 1.66 | 2.51 | 4.27 | 7.59 | 13.67 | 26.16 | 49.79 | 98.15 |

To investigate advantageous values of *t*_*max*_ for a given data set, the user of our tool has the possibility to *compute statistics* on their input data. He has to fix the following parameters beforehand: (1) The *k*-mer-size, (2) the number of hash functions and (3) the maximum false positive rate (see section 4.3.1 for sensible defaults). The statistics then give an estimation on the *expected cost* of the index size and query time of an HIBF for several values of *t*_*max*_. As an example, we computed the statistics for the real-world data of the RefSeq database from section 7.2.5 and validated them by actually building and querying a respective HIBF (Figure 7). The results show an excellent correspondence between our prediction and the actual product of run times and memory consumption, namely a correlation of 0.95 for *t*_*max*_ ranging from 64 to 8192. We were able to estimate the index size very closely to the real size, while we slightly underestimate the query time for increasing values of *t*_*max*_. In the combination of space and query time, we identified *t*_*max*_ = 192 as the optimal choice in both the expected total cost and real total cost, where 192 is a multiple of the word size 64 that is near the square root of the number of user bins *b*, i.e., 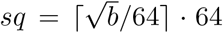. In general, our tool computes the statistics for *t*_*max*_ = *sq*, 64, 128, 256,… until the product *query time * space* increases. For the above example (Figure 7), we would stop at *t*_*max*_ = 256 since the product increases and choose *t*_*max*_ = 192 as the optimal value.

**Figure 7:**
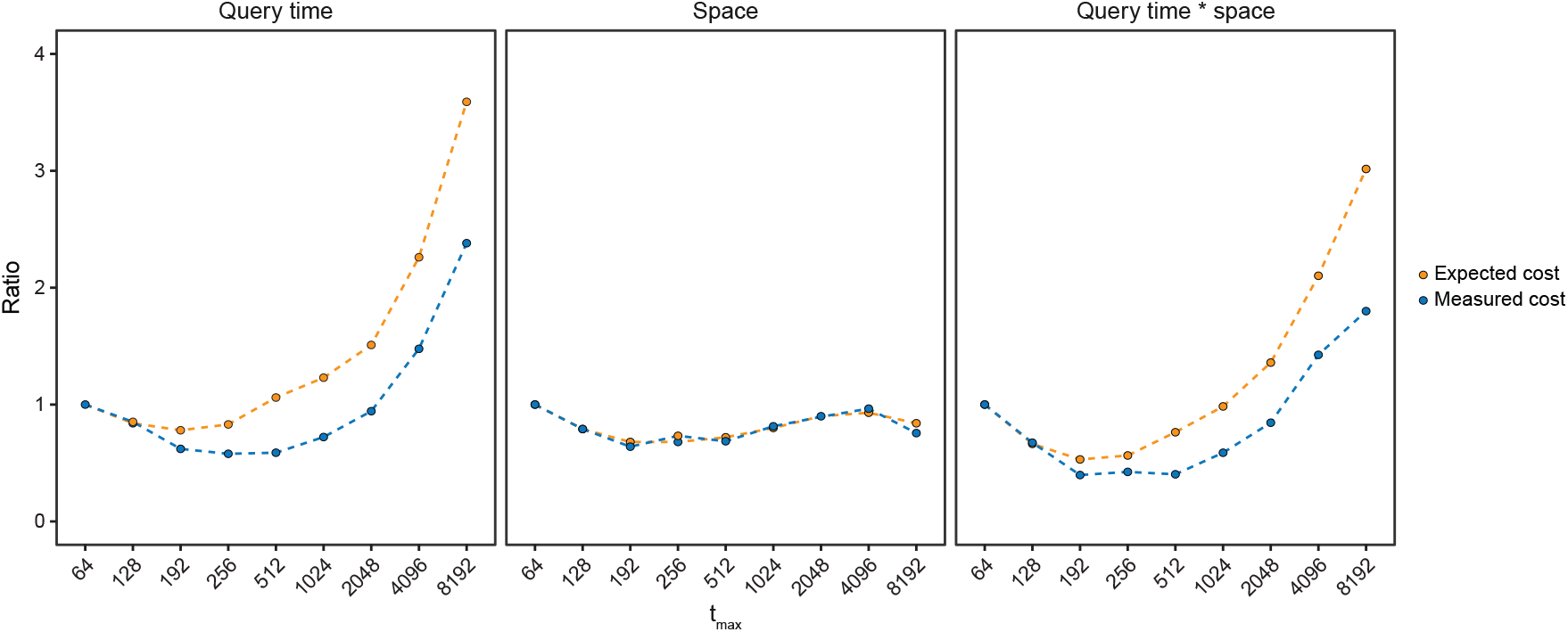
Expected cost versus real cost. The data are all complete genomes of archaea and bacteria from the RefSeq database. The relative expected cost were computed with our while the relative real cost were measured with an *t*_*max*_-HIBF with *p*_*fp*_ = 0.015, 4 hash functions and (24,20)-minimizer. The cost is given as a ratio compared to the expected/real cost of an 64-HIBF, because measurements are platform-dependent. The correlation is > 0.9, although we systematically overestimate the needed time. Nevertheless, the three best *t*_*max*_ are in both cases 192, 256 and 512 with predicted values 0.53, 0.56, 0.75 and real values 0.39, 0.42 and 0.40.

In summary, given input data, we can compute a *t*_*max*_ that minimizes the expected query time of the HIBF or the minimal expected run time weighted with memory consumption and free the user from choosing the parameter as long as the data in the UBs is not similar. We postulate that this strategy works well if a query is on average only contained in one (or a few) UBs.

#### 4.3.2 Choice of *w* and *k*

To lower the space consumption and make queries faster, we support (*w, k*)-minimizers. Other methods for shrinking the set of representative *k*-mers like described in [Lou21] or [ZKM21, Edg21] are also possible for the HIBF. For a detailed description of (*w, k*)-minimizers, see Seiler et al. [SMD^+^21]. The authors showed that for up to 2 errors that parameters *w* = 23 and *k* = 19 perform well. In general, the larger we make *w* compared to *k*, the fewer representative *k*-mers we have to store. However, the accuracy decreases, and we might increase the number of false positives and false negatives. In comparison, Kraken2 [WLL19] uses values of *w* = 35 and *k* = 31.

A *k*-mer is identified as a minimizer if it has the smallest code value in any of the windows of length *w* letters which cover it. Following the argument of Edgar [Edg21] we can define the compression factor *c*(*w, k*) to be the number of *k*-mers in a window divided by the number of minimizers, so *c*(*w, k*) ≥ 1 and larger *c*(*w, k*) indicates a smaller set of representative *k*-mers. For (*w, k*)-minimizers, *c*(*w, k*) can be estimated as follows: Consider a pair of adjacent windows of length *w* in a random sequence. Sliding one position to the right, two *k*-mers are discarded from the first window (the *k*-mer and its reverse complement) and two are added to the second. The minimizer in the second window is different from the one in the first if one of the four affected minimizers has the smallest code over both windows; otherwise, the minimizer of both windows is found in their intersection and does not change. There are 2 · (*w*−*k*+2) *k*-mers in the two windows combined, and the probability that a given one of these has the smallest code is *p* = 1/(2 · (*w* − *k* + 2)). Thus, the probability that a new minimizer is introduced by sliding the window one position is 4 · *p* = 2/(*w* − *k* + 2) and hence

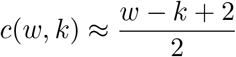

So for (23, 19)-minimizers, we would expect a compression by a factor of 3. For (32, 18)-minimizers, we would expect a compression factor of 8. So using a larger *w* is beneficial for saving space. On the other hand, the threshold for determining a match becomes smaller (roughly by the compression factor, for details see [SMD^+^21]). We want to avoid having a threshold of only 1 or 2, since then, a few false positives *k*-mer matches would result in a false positive answer.

#### 4.3.3 Impact of the HIBF’s false positive rate

Recall that we ensure that the overall false positive rate for querying an HIBF does not exceed a certain threshold. Note that, for a given false positive rate *p*_*fpr*_, the **size** *m* (bits) of an individual Bloom Filter is proportional to the amount of *k*-mers *n* it is designed to store. If *h* is the number of hash functions, then it holds:

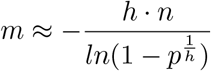

Since we aim at making each IBF in the HIBF as small as possible, we will have to accommodate a relatively high false positive rate.

We use the counts of the (*w, k*)-minimizers and the probabilistic threshold derived in [SMD^+^21] to decide whether a query is in a bin or not. However, having a relatively high false positive rate *p*_*fpr*_ will affect this thresholding. If a query has *x* minimizers, we can compute the probability that *a* of them return a false positive answer in an IBF with false positive rate *p* = *p*_*fpr*_ as

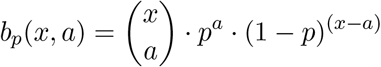

This can be quite significant. For example, for queries of length 250 we have the distribution of minimizers (computed for 1 million random reads) depicted in columns 1 and 2 of Table 4.

**Table 4:**
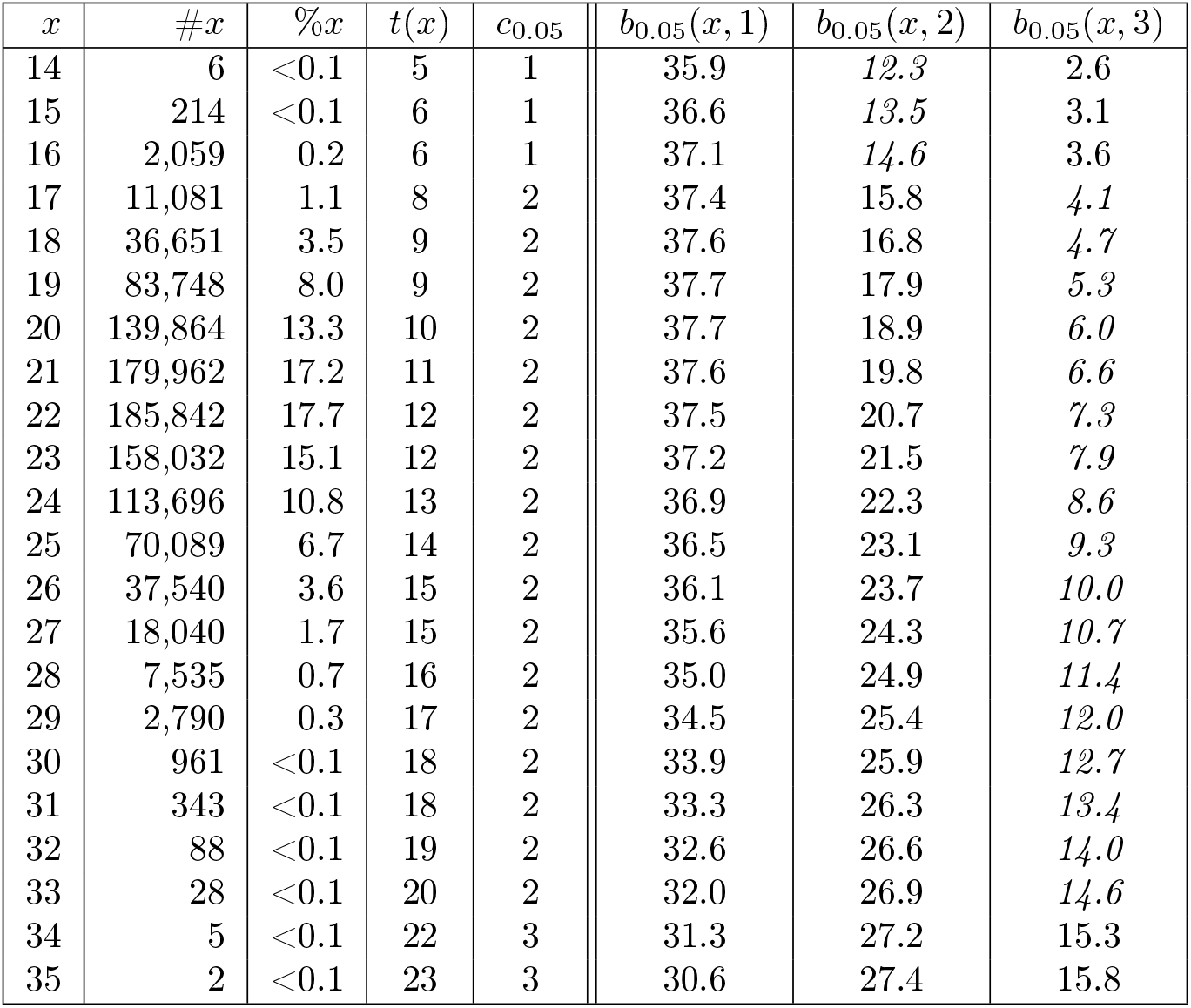
Exemplary threshold distribution. The values are for (38, 20)-minimizers, 2 errors and read length 250. Shown are the distribution #*x* (%*x*) of the number (percentage) of minimizers reaching from *x* = 14 to *x* = 35 and the threshold *t*(*x*) using the probabilistic model from [SMD^+^21]. The threshold *t*(*x*) incorporates the correction term *c*_*p*_. On the right, you see the probability *b*_*p*_(*x, a*) of having *a* false positive answers from the IBF for *p* = 0.05.

| $x$ | $\#x$ | $\%x$ | $t(x)$ | $c_{0.05}$ | $b_{0.05}(x, 1)$ | $b_{0.05}(x, 2)$ | $b_{0.05}(x, 3)$ |
| --- | --- | --- | --- | --- | --- | --- | --- |
| 14 | 6 | <0.1 | 5 | 1 | 35.9 | 12.3 | 2.6 |
| 15 | 214 | <0.1 | 6 | 1 | 36.6 | 13.5 | 3.1 |
| 16 | 2,059 | 0.2 | 6 | 1 | 37.1 | 14.6 | 3.6 |
| 17 | 11,081 | 1.1 | 8 | 2 | 37.4 | 15.8 | 4.1 |
| 18 | 36,651 | 3.5 | 9 | 2 | 37.6 | 16.8 | 4.7 |
| 19 | 83,748 | 8.0 | 9 | 2 | 37.7 | 17.9 | 5.3 |
| 20 | 139,864 | 13.3 | 10 | 2 | 37.7 | 18.9 | 6.0 |
| 21 | 179,962 | 17.2 | 11 | 2 | 37.6 | 19.8 | 6.6 |
| 22 | 185,842 | 17.7 | 12 | 2 | 37.5 | 20.7 | 7.3 |
| 23 | 158,032 | 15.1 | 12 | 2 | 37.2 | 21.5 | 7.9 |
| 24 | 113,696 | 10.8 | 13 | 2 | 36.9 | 22.3 | 8.6 |
| 25 | 70,089 | 6.7 | 14 | 2 | 36.5 | 23.1 | 9.3 |
| 26 | 37,540 | 3.6 | 15 | 2 | 36.1 | 23.7 | 10.0 |
| 27 | 18,040 | 1.7 | 15 | 2 | 35.6 | 24.3 | 10.7 |
| 28 | 7,535 | 0.7 | 16 | 2 | 35.0 | 24.9 | 11.4 |
| 29 | 2,790 | 0.3 | 17 | 2 | 34.5 | 25.4 | 12.0 |
| 30 | 961 | <0.1 | 18 | 2 | 33.9 | 25.9 | 12.7 |
| 31 | 343 | <0.1 | 18 | 2 | 33.3 | 26.3 | 13.4 |
| 32 | 88 | <0.1 | 19 | 2 | 32.6 | 26.6 | 14.0 |
| 33 | 28 | <0.1 | 20 | 2 | 32.0 | 26.9 | 14.6 |
| 34 | 5 | <0.1 | 22 | 3 | 31.3 | 27.2 | 15.3 |
| 35 | 2 | <0.1 | 23 | 3 | 30.6 | 27.4 | 15.8 |

The table shows that for a false positive rate *p*_*fpr*_ of 0.05, we encounter reads that have, e.g., 20 (38, 20)-minimizers. Those reads have a chance of almost 40 % to have one false positive count and a chance of almost 19 % to have two. This has a small effect on completely random hits. However, hits that match the query with more than the allowed errors could reach the threshold due to one or two false positive minimizers.

To counter this, we introduce a correction term for the thresholds. We add a constant *c* to the threshold *t*(*x*), which is determined by increasing *c* until the probability *b*_*p*_(*x, c*) drops below a threshold *p*_*max*_. For example, for *p* = 0.05 and *p*_*max*_ = 0.15, we have a correction of +1 for 14 ≤ *x* ≤ 16 and a correction of +2 for 17 ≤ *x* ≤ 33. For *p* = 0.02, we have a correction of +1 for all *x*. The value for *p*_*max*_ was experimentally determined using the benchmark from [SMD^+^21] such that the benchmark yielded no false positives and no false negatives. It is set to *p*_*max*_ = 0.15. Those corrections are precomputed and incorporated into the thresholding.

### 4.4 Recommended choice of parameters

As pointed out above, we want to free the average user from setting internal parameters of the HIBF. Here we describe how the HIBF will set its internal key parameters depending on a few input parameters provided by the user: 1) the sequences in user bins, 2) the length of the query sequences (e.g., 150 or 250 long Illumina reads or high quality PacBio reads of several thousand length), and 3) the maximum number of allowed errors. Then the HIBF will proceed as follows:

1. Choice of *h* and *p*_*fpr*_ The choice of the number of hash functions *h* influences the false positive rate, depending on the allocated space for the HIBF. In our experiments, the values of *h* =4 and *p*_*fpr*_ = 0.015 turned out to be suitable for all analyzed data sets.
2. Choice of *k* Depending on the error rate, we will choose *k* such that the *k*-mer lemma has still a positive threshold. For example, when querying reads of length 200 and allowing 4 errors, the error rate is 2% and therefore *k* has to be less than 67. We will choose *k* such that a random *k*-mer match in the database is unlikely. For example, for 100 GiB RefSeq data, we would choose *k* = 20.
3. Choice of *w* Depending on the choice of *k*, the choice of *w* has to be made, such that we obtain positive thresholds (in general, we aim at having a minimum threshold of 3 for the (*w, k*) minimizers. Hence, we choose *w* as large as possible, such that the minimum threshold is 3. For example, this is obtained for 2 errors and read length 150 for (29, 20) minimizers, for read length 100 for (24, 20)-minimizers and for read length 250 for (40, 20) minimizers, which in turn reduces the amount of *k*-mers by factors of 5.5, 3, and 11, respectively.
4. Choice of *t*_*max*_ After the choice of (*w, k*) minimizers, we compute *t*_*max*_ as pointed out in Section 4.3.1 and then build the HIBF with a false positive rate of 0.015, a value that has proven to be a good compromise between space consumption and needed threshold correction. When we query, we use our probabilistically derived threshold, which we correct depending on the false positive rate of the IBF as described above.

## 5 Data availability

All scripts and supplemental code used in this paper are available at https://github.com/seqan/raptor. Scripts include instructions for re-simulating the data sets of section 2.2.1. The RefSeq database as of January 28th 2022 was downloaded via the *genome_updater*. The RNA-Seq data set accessions are available at https://doi.org/10.5281/zenodo.1186393.

## 6 Software and Code

*Chopper* (computes the HIBF layout) and *Raptor* are written in C++20 using the SeqAn Library [RDE^+^17] and available at https://github.com/seqan/chopper and https://github.com/seqan/raptor, respectively. All software is listed in the following table:

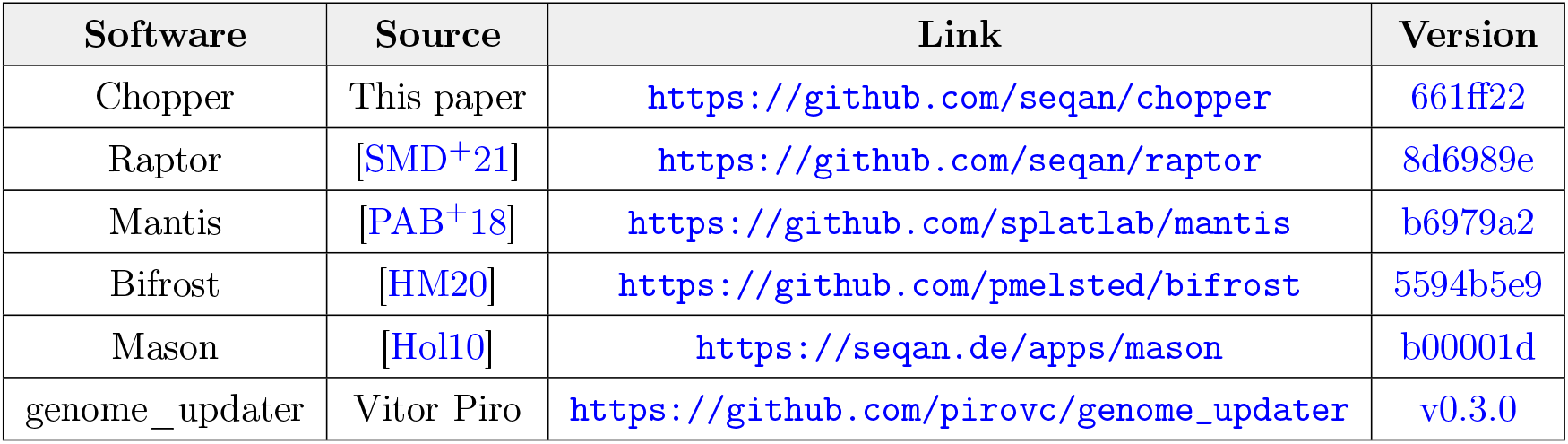

| Software | Source | Link | Version |
| --- | --- | --- | --- |
| Chopper | This paper | https://github.com/seqan/chopper | 661ff22 |
| Raptor | [SMD<sup>+</sup>21] | https://github.com/seqan/raptor | 8d6989e |
| Mantis | [PAB<sup>+</sup>18] | https://github.com/splatlab/mantis | b6979a2 |
| Bifrost | [HM20] | https://github.com/pmelsted/bifrost | 5594b5e9 |
| Mason | [Hol10] | https://seqan.de/apps/mason | b00001d |
| genome_updater | Vitor Piro | https://github.com/pirovc/genome_updater | v0.3.0 |

## Footnotes

1 https://github.com/Cyan4973/xxHash/tree/v0.7.3

2 https://github.com/hideo55/cpp-HyperLogLog/tree/517598b2

