## Supplementary material for "Hierarchical Interleaved Bloom Filter: Enabling ultrafast, approximate sequence queries": HIBF_supplement.pdf

### 7 Supplement

#### 7.1 Hardware specifications

- Platform: Dell PowerEdge T640
- CPU:
  - Intel Xeon Gold 6248
  - 2 Sockets, 20 Cores per Socket, 2 Threads per Core
  - 40 physical CPUs, 80 logical CPUs (threads)
  - 3.9 GHz
- RAM:
  - 1 TiB
  - 2,666 MHz
  - 16 · HMAA8GR7AJR4N-WM (64 GiB each)

#### 7.2 Results

| | **Pre-build** | | **Build** | | | **Query** | | **Statistics** | |
|---|---|---|---|---|---|---|---|---|---|
| **Method** | Time | RAM | Time | RAM | Index Size | Time | RAM | FP | FN |
| **20-mer** | | | | | | | | | |
| IBF 5% | none | | 7:59 | **4.3 GiB** | **4.1 GiB** | 511:53 | **7.6 GiB** | 0 | 0 |
| IBF 1.5% | none | | 8:09 | 8.3 GiB | 8.1 GiB | 489:39 | 11.6 GiB | 0 | 0 |
| 1024-HIBF 5% | 1788:18 | 32.7 GiB | **1:08** | 10.5 GiB | 8.0 GiB | 1:56 | 11.4 GiB | 0 | 0 |
| 1024-HIBF 1.5% | 1788:31 | 32.7 GiB | 1:57 | 18.3 GiB | 16.0 GiB | **1:47** | 18.8 GiB | 0 | 0 |
| **(24,20)-minimizer, tau 0.99, pmax 0.4** | | | | | | | | | |
| IBF 5% | none | | 2:42 | **1.9 GiB** | **1.7 GiB** | 175:35 | **5.1 GiB** | 0 | 0 |
| IBF 1.5% | none | | 2:10 | 3.3 GiB | 3.1 GiB | 129:45 | 6.6 GiB | 0 | 0 |
| 1024-HIBF 5% | 1841:42 | 32.7 GiB | **0:27** | 4.3 GiB | 2.7 GiB | 0:57 | 6.2 GiB | 0 | 0 |
| 1024-HIBF 1.5% | 1814:06 | 32.7 GiB | 0:29 | 6.6 GiB | 5.1 GiB | **0:52** | 8.5 GiB | 0 | 0 |

Table 5: **Benchmark results on simulated data of** $1,048,576$ ***user bins***. The data compose 64 GiB of random DNA sequence split into $1,048,576$ *user bins* and one million queries of length 250 bp generated from the former. *Pre-build* includes all steps that have to be done before the actual index is built (online methods section 4.2.1). Both Bifrost and Mantis crash when building the index.

All measurements of the one million user bin simulation can be found in table 5. All measurements used to produce figure 4 can be found in table 6.

##### 7.2.1 Simulated 65,536 result file sizes

Due to different output formats, the output sizes of *Bifrost*, *Mantis*, and *Raptor* differ dramatically. Besides the occupied disk space, this may also have implications for downstream analyses. All three programs report their results as plain text files and report the full filename of the bins. *Bifrost* outputs a matrix, where each column is a bin and each row is a read ID. A 0 or 1 in a cell indicates the absence or presence of a read in a bin, respectively. Searching 10 million reads in $65,536$ bins results in an output file of size 1.3 TiB. *Mantis* lists for each read the $k$-mer counts for all bins in which $k$-mers were found.

| **Method** | **Pre-build** Time | **Pre-build** RAM | **Build** Time | **Build** RAM | **Build** Index Size | **Query** Time | **Query** RAM | **Statistics** FP | **Statistics** FN | **Statistics** Accuracy |
|---|---|---|---|---|---|---|---|---|---|---|
| **20-mers** | | | | | | | | | | |
| 192-HIBF 1.5% | 13:20 | 2.38 GiB | 36:23 | 214.8 GiB | 159.15 GiB | 5:32 | 162.98 GiB | 0.022 % | 0 | 99.977 % |
| **24-mers** | | | | | | | | | | |
| 192-HIBF 1.5% | 13:13 | 2.36 GiB | 38:32 | 230.07 GiB | 160.78 GiB | 4:51 | 164.61 GiB | 0.030 % | 0 | 99.969 % |
| **28-mers** | | | | | | | | | | |
| 192-HIBF 1.5% | 13:01 | 2.35 GiB | 42:01 | 238.26 GiB | 162.02 GiB | 4:59 | 165.85 GiB | 0.039 % | 0 | 99.960 % |
| **32-mers** | | | | | | | | | | |
| Mantis | 140:35 | 1.4 GiB | 2719:19 | 796.4 GiB | 468.0 GiB | 131:54 | 496.4 GiB | **0.0001** % | 0.04851 % | 99.951 % |
| IBF 5% | none | | **32:37** | 293.1 GiB | 292.5 GiB | 13:08 | 296.3 GiB | 0.048 % | **0** | 99.951 % |
| IBF 1.5% | none | | 40:02 | 435.4 GiB | 343.7 GiB | 16:14 | 438.6 GiB | **0.047 %** | **0** | 99.952 % |
| 192-HIBF 5% | 12:58 | 2.4 GiB | 54:13 | 210.9 GiB | **109.9 GiB** | **4:55** | 113.8 GiB | 0.058 % | **0** | 99.941 % |
| 192-HIBF 1.5% | 12:53 | 2.4 GiB | 60:27 | 253.4 GiB | 163.1 GiB | 5:23 | 167.0 GiB | 0.051 % | **0** | 99.949 % |
| **(24,20)-minimizer, tau 0.99, pmax 0.4** | | | | | | | | | | |
| Bifrost | none | | 1040:27 | 519.7 GiB | 265.3 GiB | 232:54 | 575.7 GiB | 0.905 % | 0.3239 % | 98.770 % |
| IBF 5% | none | | **11:24** | 93.0 GiB | 92.3 GiB | 4:19 | 96.2 GiB | 0.113 % | 0.0003 % | 99.886 % |
| IBF 1.5% | none | | 12:01 | 137.9 GiB | 137.3 GiB | 6:23 | 141.1 GiB | **0.112 %** | **0.0003 %** | 99.887 % |
| 192-HIBF 5% | 13:36 | 2.4 GiB | 12:55 | 57.23 GiB | **34.4 GiB** | **1:48** | 38.3 GiB | 0.128 % | 0.0003 % | 99.871 % |
| 192-HIBF 1.5% | 13:23 | 2.4 GiB | 15:23 | 78.37 GiB | 51.1 GiB | 1:50 | 54.9 GiB | 0.116 % | 0.0003 % | 99.882 % |
| **(32,20)-minimizer, tau 0.99, pmax 0.4** | | | | | | | | | | |
| 192-HIBF 1.5% | 13:28 | 2.36 GiB | 5:39 | 31.43 GiB | 22.14 GiB | 1:04 | 25.97 GiB | 0.302 % | 0.00003 % | 99.698 % |
| **(40,20)-minimizer, tau 0.99, pmax 0.4** | | | | | | | | | | |
| 192-HIBF 5% | 12:55 | 2.35 GiB | 3:56 | 15.80 GiB | **9.62 GiB** | 2:49 | 13.45 GiB | 0.465 % | 0.00001 % | 99.534 % |
| 192-HIBF 1.5% | 13:34 | 2.36 GiB | 4:04 | 20.66 GiB | 15.11 GiB | **1:07** | 18.94 GiB | 0.452 % | 0.00001 % | 99.547 % |
| **(28,24)-minimizer, tau 0.99, pmax 0.4** | | | | | | | | | | |
| 192-HIBF 1.5% | 13:10 | 2.36 GiB | 12:12 | 73.88 GiB | 51.59 GiB | 1:31 | 55.42 GiB | 0.115 % | 0.0004 % | 99.884 % |
| **(32,24)-minimizer, tau 0.99, pmax 0.4** | | | | | | | | | | |
| 192-HIBF 1.5% | 13:08 | 2.35 GiB | 7:31 | 43.01 GiB | 30.87 GiB | 1:11 | 34.69 GiB | 0.229 % | 0.00003 % | 99.770 % |
| **(40,32)-minimizer, tau 0.99, pmax 0.4** | | | | | | | | | | |
| 192-HIBF 1.5% | 13:06 | 2.36 GiB | 7:28 | 35.32 GiB | 31.49 GiB | 1:09 | 35.32 GiB | 0.242 % | 0.00001 % | 99.758 % |

Table 6: **All complete genomes of Archaea and Bacteria in RefSeq**. The uncompressed data set has a size of about 98.8 GiB. Query reads of length 250 *bp* were simulated using the *Mason simulator* [Hol10]. *Mantis* does not support minimizers and could only be used with $k = 32$ because it crashed for $k = 20$. Although *Mantis* is technically an exact method outputting $k$-mer counts, a threshold (in this case 0.7) needs to be applied to determine the query membership resulting in few false positives/negatives. Bifrost was run with $(24, 20)$-minimizers and a threshold of 0.36. For details on the chosen thresholds, see section 2.2.

The resulting file is 1.1 GiB big, roughly 1,200 times smaller than *Bifrost*'s output. *Raptor* assigns each bin a number inside a header section, and then provides for each read ID a list of matching bins. The output file is 132 MiB big. This is a million times smaller than *Bifrost* and 8 times smaller than *Mantis*.

##### 7.2.2 RNA-Seq data

In addition to the artificial data and the RefSeq benchmark, we also performed some benchmarks on RNA-Seq data.

This real-world data set consists of 1,742 RNA-Seq files and is a subset of the data used as a benchmark in similar applications [PAB+18]. Similar to [SMD+21], experiments with a read length below 50 bp were excluded. Low frequency $k$-mers, which are likely the result of sequencing errors, are excluded by applying the same cutoffs as [PAB+18]. Unlike previous analysis, this analysis was performed with one thread to stay consist with previous analysis [PAB+18, SMD+21]. Since the benefits of the HIBF over the IBF were shown in

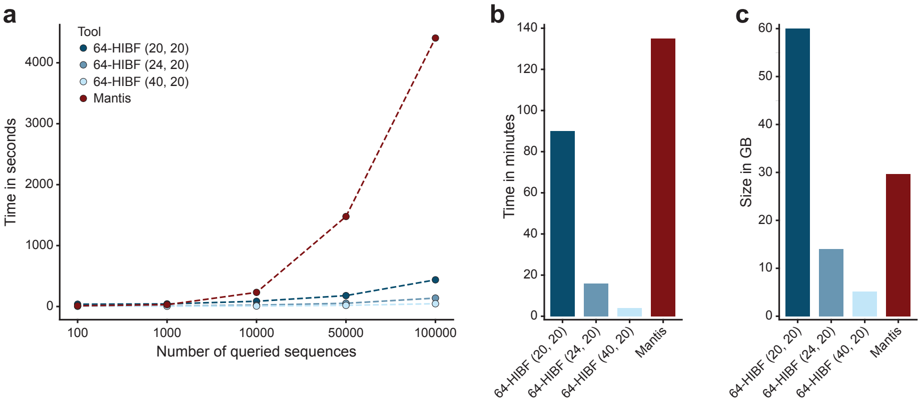


Figure 8: **RNA-Seq results**. (a) The query time for the HIBF and Mantis for 100/1,000/10,000/ 50,000/100,000 sequences with a threshold of 0.8. Times are in seconds. (b) The index sizes in GiB and the built-times in minutes.

the previous sections, we focus on the comparison of the HIBF against its competitor *Mantis*.

For this data set, the HIBF can be built faster than Mantis for any settings. The resulting index is larger than that of Mantis for $k$-mers and less for $(24, 20)$-minimizers and $(40, 20)$-minimizers. Indeed, for $(40, 20)$-minimizers less than a third compared to Mantis (see Figure 8a).

Furthermore, we compared the query performance of Mantis and the HIBF by searching for $100/1,000/10,000/50,000/100,000$ random transcripts from the human genome with a threshold of 0.8. As can be seen in Figure 8b, the HIBF outperforms Mantis by orders of magnitude for all built HIBFs once the number of queries is greater than $1,000$. With fewer queries, only the HIBFs with $(24, 20)$-minimizers and $(40, 20)$-minimizers are faster than Mantis. We believe that the query times in the beginning are heavily influenced by loading the index, which is the reason the HIBF with $k$-mers is a bit slower than Mantis.

##### 7.2.3 Data simulation of 65,536 user bins

Following the approach in [SMD+21] we created a random DNA sequence of 4 GiB size and divided it into $b$ bins which would correspond to $b$ different genomes, e.g., a metagenomic data set. Using the Mason genome variator [Hol10], we then generated 16 similar genomes in each bin, which differ about 1% from each other on average. This could be seen as bins containing the genomes for a homologous species. The total sequence length is hence 64 GiB. Finally, we uniformly sampled one million $(2^{2}0)$ reads of length 250 bp from the genomes and introduced 2 errors in each read to simulate a sequencing experiment. This artificial data set is well-balanced, which is the ideal case for the IBF, since its overall size is dependent on the largest bin. What we change is that we simulated many more bins, namely $b = 2^{16} = 65,536$ and $b = 2^{20} = 1,048,576$ to show the detrimental effect of many user bins to the IBF and how the HIBF removes the limitation.

##### 7.2.4 The RefSeq data set

The data set contained all RefSeq Archaea and Bacteria Complete Genomes as of 28-01-2022, with a size of 98.8 GiB uncompressed and 28.5 GiB compressed. The sequences were in FASTA format. The data set has 25,321 files and 54,348 FASTA records.

##### 7.2.5 Statistics of different values for $t_{max}$ for the RefSeq data set

We computed statistics with the tool *Chopper* on this data set to choose a fitting $t_{max}$. The results are shown in table 7 (*query cost, space cost* and *total cost*). We validated the statistics by actually building and querying a respective HIBF (*real time, real mem* and *real total*).

| $t_{max}$ | **64** | **128** | **sq=192** | **256** | **512** | **1024** | **2048** | **4096** | **8192** |
|---|---|---|---|---|---|---|---|---|---|
| *bin penalty* | 1.00 | 1.05 | 1.29 | 1.67 | 2.37 | 3.34 | 5.80 | 9.63 | 17.59 |
| *query cost* | 1.00 | 0.84 | **0.78** | 0.83 | 1.06 | 1.23 | 1.51 | 2.26 | 3.59 |
| *space cost* | 1.00 | 0.79 | **0.68** | **0.68** | 0.72 | 0.80 | 0.90 | 0.93 | 0.84 |
| total cost | 1.00 | 0.66 | **0.53** | 0.56 | 0.76 | 0.98 | 1.36 | 2.10 | 3.01 |
| real time | 1.00 | 0.85 | 0.62 | 0.58 | 0.59 | 0.72 | 0.94 | 1.48 | 2.38 |
| real mem | 1.00 | 0.79 | 0.64 | 0.73 | 0.69 | 0.81 | 0.90 | 0.97 | 0.76 |
| real total | 1.00 | 0.67 | 0.39 | 0.42 | 0.40 | 0.59 | 0.84 | 1.43 | 1.80 |

Table 7: **Expected relative runtimes, memory consumption and the runtime relative to memory consumption for different choices of $\mathbf{t_{max}}$ and $\mathbf{p_{fpr} = 0.0125}$.** *bin penalty* refers to the increase in runtime when querying an original IBF with $t_{max}$ bins. All number are given as ratios compared to the (H)IBF version with 64 (user) bins. *query, space* and *total cost* refers are estimated cost of an $t_{max}$-HIBF computed by the tool *Chopper*. *real time, real mem* and *real total* refer to the experimentally derived ratios of an $t_{max}$-HIBF using (24,20)-minimizer, 4 hash functions and an $p_{fpr} = 0.0125$. The minimum time is reached for $t_{max} = 192$ . The minimum relative run time is also reached for $t_{max} = 192$, followed by the value for $t_{max} = 256$.
